# Kinetic Plasticity of Nitrite-Oxidizing Bacteria Containing Cytoplasmic Nitrite Oxidoreductase

**DOI:** 10.1101/2025.07.31.663499

**Authors:** Ui-Ju Lee, Joo-Han Gwak, Christiana Abiola, Seongjun Lee, Jin-Sun Yu, Ok-Ja Si, Hyo Je Cho, Zhe-Xue Quan, Katharina Kitzinger, Holger Daims, Michael Wagner, Man-Young Jung, Sung-Keun Rhee

**Author notes:** Corresponding authors. (S-K R) & (M-Y J).

## Abstract

Nitrite oxidation, the second step of nitrification, is essential to the global nitrogen cycle. Nitrite-oxidizing bacteria (NOB) are classified into two groups based on the cellular localization of their key enzyme nitrite oxidoreductase (NXR): periplasmic (pNXR) and cytoplasmic (cNXR). The use of a cNXR by NOB has been reported to be linked to a lower nitrite affinity and energy efficiency of nitrite oxidation, indicating adaptation to nitrogen-rich environments. In this study, cNXR NOB model strains demonstrated nitrite concentration-dependent shifts in optimal growth pH, a behavior not observed in pNXR NOB. *Nitrobacter winogradskyi* Nb-255 (cNXR NOB), grown at 1 mM nitrite (pH 7.5), exhibited a high nitrite affinity in terms of apparent *K*_m_ (25.9 μM) and a high specific affinity *a°* (440.5 l g cells^−1^ h^−1^), both comparable to pNXR NOB in microrespirometry-based kinetic assays. Unexpectedly, cells pre-grown at 10 mM nitrite (pH 7.5) achieved a pNXR-like affinity at pH 5.5 without prior adaptation to acidic conditions. In contrast, pNXR NOB exhibited consistent kinetic behavior across different pH conditions. Kinetic inhibition in the presence of nitrate suggested that this plasticity is driven by a regulated interplay between nitrite uniport and nitrite/nitrate antiporter systems. Our findings indicate that *Nitrobacter* can dynamically modulate nitrite affinity in response to both nitrite concentration and pH, conferring a flexible adaptation strategy that features traits of both *r*-and *K*-strategists across a range of environmental conditions. This adaptive plasticity likely extends to other cNXR-containing NOB in response to fluctuating environmental conditions.

## Introduction

Nitrite-oxidizing bacteria (NOB) catalyze the conversion of nitrite (NO_2_⁻) to nitrate (NO_3_⁻), the second step of nitrification, a crucial process in the global biogeochemical nitrogen cycle^1^. Despite their pivotal role across diverse ecosystems, NOB have been understudied compared to other microorganisms involved in the nitrogen cycle, partly due to the prevailing assumption that they depend on ammonia-oxidizing microorganisms (AOM) for their nitrite supply^1^. However, this perspective oversimplifies NOB physiology and neglects the critical, independent functions of NOB in diverse ecosystems.

Nitrite is a toxic and reactive intermediate^2^ that must be rapidly removed to prevent its accumulation and the self-intoxication of AOM^3,4^. The ecological dynamics of NOB are tightly linked to their interactions with other microbial groups and nutrient fluxes in diverse environments^5,6^. Emerging evidence reveals that NOB are more metabolically versatile than previously assumed. In addition to oxidizing nitrite, members of some NOB lineages have been shown to utilize alternative electron donors such as hydrogen^7–9^ and formate^6,10^, reduce nitrate under low-oxygen conditions^6,9,11^, and engage in reciprocal feeding with AOMs^5,6^. Building on this metabolic flexibility, understanding how diverse NOB taxa adapt to environmental perturbations across ecosystems is increasingly critical for elucidating their ecological functions and contributions to nitrification.

Known NOB belong to twelve genera from four bacterial phyla: *Nitrobacter*^12^, ‘*Candidatus*. Nitrotoga’^13^, and *Nitrococcus*^14^ (all *Pseudomonadota*); *Nitrospira*^9^ and ‘*Ca*. Nitronereus’^15^ (both *Nitrospirota*); *Nitrospina*^16^, ‘*Ca.* Nitromaritima’^17^, ‘*Ca*. Nitrohelix’^18^, and ‘*Ca*. Nitronauta’^18^ (all *Nitrospinota*); *Nitrolancea*^19^, ‘*Ca*. Nitrocaldera’^20^, and ‘*Ca*. Nitrotheca’^20^ (all *Chloroflexota*). The key enzyme driving nitrite oxidation in NOB is nitrite oxidoreductase (NXR), which catalyzes the conversion of nitrite to nitrate while transferring electrons into the respiratory chain. NXR typically consists of three subunits – NxrA (α), NxrB (β), and NxrC (γ) – and its cellular localization varies among NOB genera^1^. In all *Nitrospirota* and *Nitrospinota* NOB genera, as well as in ‘*Ca*. Nitrotoga’, and ‘*Ca.* Nitrotehca’, NxrA is located in the periplasmic space (with the NXR holoenzyme either being membrane-bound, or entirely periplasmatic), whereas in all other known NOB, NXR is membrane-bound with the active site facing the cytoplasm. A notable exception is ‘*Ca*. Nitrocaldera’ which has both cytoplasmic-facing and periplasmic NXR forms^20^.

The subcellular localization of NXR influences nitrite affinity and energy efficiency during nitrite oxidation^1,21^. Cytoplasmic NXR (cNXR) requires the transport of nitrite and nitrate across the cytoplasmic membrane, which may create a physiological bottleneck depending on the efficiency of the transporter. Additionally, in cNXR systems, protons released from water during nitrite oxidation within the cytoplasm do not directly contribute to the proton motive force, leading to reduced cellular energy yields compared to periplasmic NXR (pNXR) systems^21^. These differences probably play a significant role in the niche differentiation between cNXR-containing genera (e.g., *Nitrobacter*), which thrive in nitrogen-rich conditions^22–24^, and pNXR-containing genera (e.g., *Nitrospira*), which dominate in low-nitrite environments^22,23,25^.

Nitrification also occurs even in acidic soils^26–28^, which are widespread across the Earth’s terrestrial ecosystems^29^; however, nitrite concentrations in these environments are typically very low or even undetectable^30^. This is possibly caused by high nitrite oxidation rates mediated by NOB^31–33^, abiotic decomposition of nitrite under acidic conditions via the formation and instability of nitrous acid (HNO_2_)^34,35^, and atmospheric release of HNO_2_ as gas-phase HONO^30^. Surprisingly, members of the genus *Nitrobacter*, which possess cNXR with relatively high predicted *K*_m_ values (indicating low substrate affinity)^36^, have been reported as numerically dominant in acidic soils with pH ranges as low as 4.3 to 5.2^37^. These observations contradict the established assumptions about the ecological niche of *Nitrobacter* as a copiotroph adapted to neutral pH environments characterized by elevated nitrite concentrations^22^.

In this study, we conducted physiological and kinetic analyses of an acid-adapted *Nitrobacter* strain isolated from forest soil to explore its environmental adaptations. Further, the study was expanded to include diverse representative cNXR and pNXR NOB for whole-cell-based comparative kinetic analysis. Unexpectedly, cNXR NOB exhibited dynamic shifts in nitrite affinity depending on nitrite concentrations and pH. These findings offer new insights into the adaptive plasticity of cNXR NOB, which were previously considered kinetically-restricted copiotrophs.

## Results and Discussion

### Isolation of an Acid-Adapted Nitrite-Oxidizing Bacterium

A nitrite-oxidizing bacterium was enriched after inoculating mineral NOB medium with acidic forest soil from Jeju Island (Supplementary Table 1) and incubated at pH 5.0 (resembling the pH of the original soil sample) using 100 μM nitrite as the sole energy and nitrogen source. After multiple rounds of subculturing and serial dilutions of this enrichment culture in fresh medium, a pure *Nitrobacter* strain, designated as ‘strain JJSN’, was successfully isolated. Strain JJSN exhibited exponential growth in batch culture, as indicated by an increase in 16S rRNA gene copy number, with a nearly stoichiometric conversion of nitrite to nitrate (Fig. 1a). Whole-genome sequencing of strain JJSN was performed (Supplementary Table 2; Supplementary Dataset 1), and phylogenomic analysis based on conserved concatenated marker protein sequences confirmed that strain JJSN belongs to the *Nitrobacter* genus (Fig. 1b). Based on average nucleotide identity (ANI) (Supplementary Fig. 1a) and average amino acid identity (AAI) (Supplementary Fig. 1b) analyses, strain JJSN appears to represent a distinct novel species within the genus *Nitrobacter*. The NxrA of strain JJSN showed the highest amino acid sequence identity (95.2%) with those of *Nitrobacter* strains (ECS31B4, HKST-UBA79, and Nb-311A) isolated from diverse environments (Supplementary Fig. 1c).

**Fig. 1:**
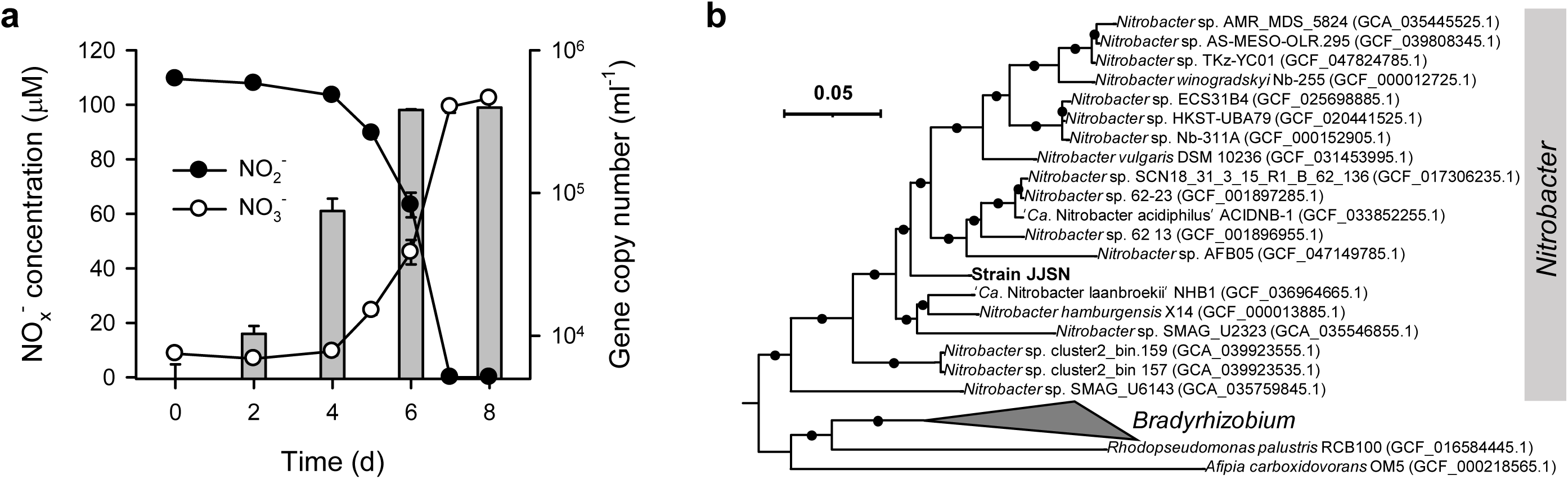
Growth curve and phylogenomic tree of the acidophilic nitrite-oxidizing bacterium, strain JJSN. **a**, Growth curve of strain JJSN showing nitrite oxidation, nitrate production, and cell abundance of strain JJSN over time. Samples were collected every two days; the bar represents 16S rRNA gene copy number. Data are presented as mean ± s.d. (*n* = 3). **b,** Phylogenomic tree of *Nitrobacter* strains inferred based on the concatenated alignment of 71 phylogenetic marker genes across 28 genomes and metagenome-assembled genomes (MAGs) retrieved from the NCBI database. Only genomes and MAGs with ≥ 50% completeness and ≤ 5% contamination were included. Branch support values ≥ 95% are indicated by black circles.

### Adaptation to Acidic Conditions

The effect of pH on the nitrite oxidation activity was examined using batch cultures of strain JJSN across four different initial nitrite concentrations (10, 100, 1000, and 10,000 μM) (Fig. 2). At pH 4.0, there was significant abiotic nitrite loss without any corresponding production of nitrate (Supplementary Fig. 2), aligning with previous studies^34,35^. As a result, subsequent experiments were conducted at pH levels ranging from 4.5 to 9.0 for strain JJSN. At lower nitrite concentrations (10 and 100 μM), strain JJSN exhibited nitrite oxidation activity within a pH range of 4.5 to 6.0, with the highest oxidation rate observed at pH 5.0 (Fig. 2a; Supplementary Figs. 3a–c), highlighting its acidophilic characteristics. Conversely, at higher nitrite concentrations (1,000 and 10,000 μM), the optimal pH shifted to a range of 6.0 to 7.0 (Fig. 2a), and no nitrite oxidation activity was detected below pH 6.0. This shift toward a more acidic optimal pH at low nitrite concentrations likely confers ecological advantages on strain JJSN in acidic terrestrial environments, where nitrite availability is limited and HNO_2_ toxicity is high^38^.

**Fig. 2:**
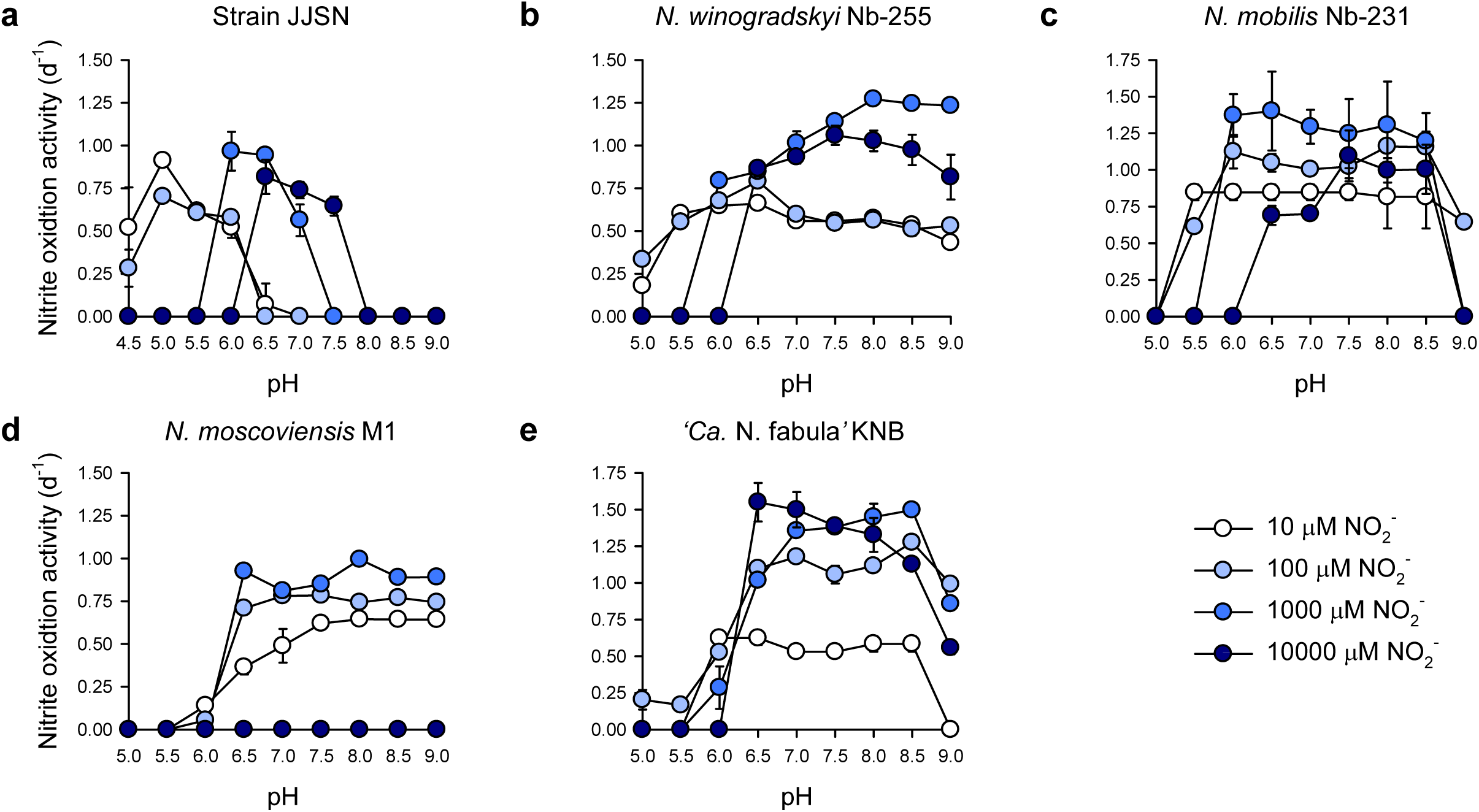
Nitrite oxidation activity of NOB strains across varying pH and nitrite concentrations. Nitrite oxidation activity was measured in five strains: **a**, Strain JJSN; **b**, *N. winogradskyi* Nb-255; **c**, *N. mobilis* Nb-231; **d**, *N. moscoviensis* M1; and **e**, ‘*Ca*. N. fabula’ KNB. Nitrite oxidation activity during the exponential phase in media supplemented with four nitrite concentrations: 10 µM (white), 100 µM (light blue), 1,000 µM (medium blue), and 10,000 µM (dark blue), with initial pH ranging from 5.0 to 9.0 (4.5 to 9.0 for strain JJSN). Data represent mean ± s.d. (*n* = 3). Nitrite oxidation profiles used for rate calculations are provided in **Supplementary** Fig. 3.

For comparison, a neutrophilic nitrite-oxidizing bacterium, *Nitrobacter winogradskyi* Nb-255, was tested under the same conditions. At lower nitrite concentrations (10 and 100 μM), *N. winogradskyi* Nb-255 demonstrated nitrite oxidation over a broader pH range of 5.0 to 9.0, without a distinct optimal pH (Fig. 2b; Supplementary Figs. 3e–h). However, at higher nitrite concentrations (1,000 and 10,000 μM), the optimal pH shifted to a range of 7.5 to 9.0, with an increase in nitrite oxidation rates, reflecting its neutrophilic characteristics (Fig. 2b; Supplementary Figs. 3g, h). Interestingly, both *Nitrobacter* strains exhibited shifts in their optimal pH for nitrite oxidation in response to the nitrite concentrations, regardless of whether they were acidophilic or neutrophilic. Recently, variation in the pH responses at different nitrite concentrations has also been observed in ‘*Ca.* Nitrobacter laanbroekii’ NHB1^33^.

To determine whether the shift in optimal pH for nitrite oxidation activity in response to different nitrite concentrations is specific to the genus *Nitrobacter*, we tested other representative cultivated NOB, including cNXR-containing (*Nitrococcus mobilis* Nb-231) and pNXR-containing (*Nitrospira moscoviensis* M1 and ‘*Ca.* Nitrotoga fabula’ KNB) strains (Figs. 2c–e; Supplementary Figs. 3i–s). As observed in *N. winogradskyi* Nb-255 (see Fig. 2b), *N. mobilis* Nb-231 retained nitrite oxidation activity at lower nitrite concentrations (10 and 100 μM) across a similarly wide pH range of 5.5 to 9.0 (Fig. 2c; Supplementary Figs. 3i–l). In contrast, neither *N. moscoviensis* (Figs. 2d; Supplementary Figs. 3m–o) nor ‘*Ca.* N. fabula’ KNB (Fig. 2e; Supplementary Figs. 3p–s) showed variation in their optimal pH for nitrite oxidation activity across different nitrite concentrations. Taken together, these findings suggest that the dynamic shifts in optimal pH in response to varying substrate (nitrite) concentrations during cultivation are distinct features of cNXR NOB, such as *Nitrobacter* and *Nitrococcus,* but not of pNXR NOB. This apparent flexibility possibly reflects their unique adaptive ecological strategies for thriving in environments with fluctuating pH and nitrogen availability.

### Kinetic Analysis

#### Effect of the Growth Nitrite Concentration on Affinity

The variation in optimal pH for nitrite oxidation observed in cNXR NOB may be attributed to differences in nitrite affinity, which is influenced by the initial nitrite concentrations used during cultivation. As nitrite toxicity increases under acidic conditions due to its protonation to HNO_239_, a high-affinity nitrite oxidation system may help protect cells from this acid stress. To investigate this, we compared the nitrite affinity of *N. winogradskyi* Nb-255 cells grown at 1 mM and 10 mM nitrite at pH 7.5 using microrespirometry (MR)-based whole-cell kinetic analysis (Fig. 3a, b). The kinetics of nitrite oxidation, measured via oxygen uptake (Eq. 1), followed the expected stoichiometry of the reaction (Supplementary Fig. 4) and were consistent with previous studies^36^:

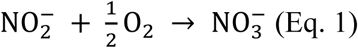

**Fig. 3:**
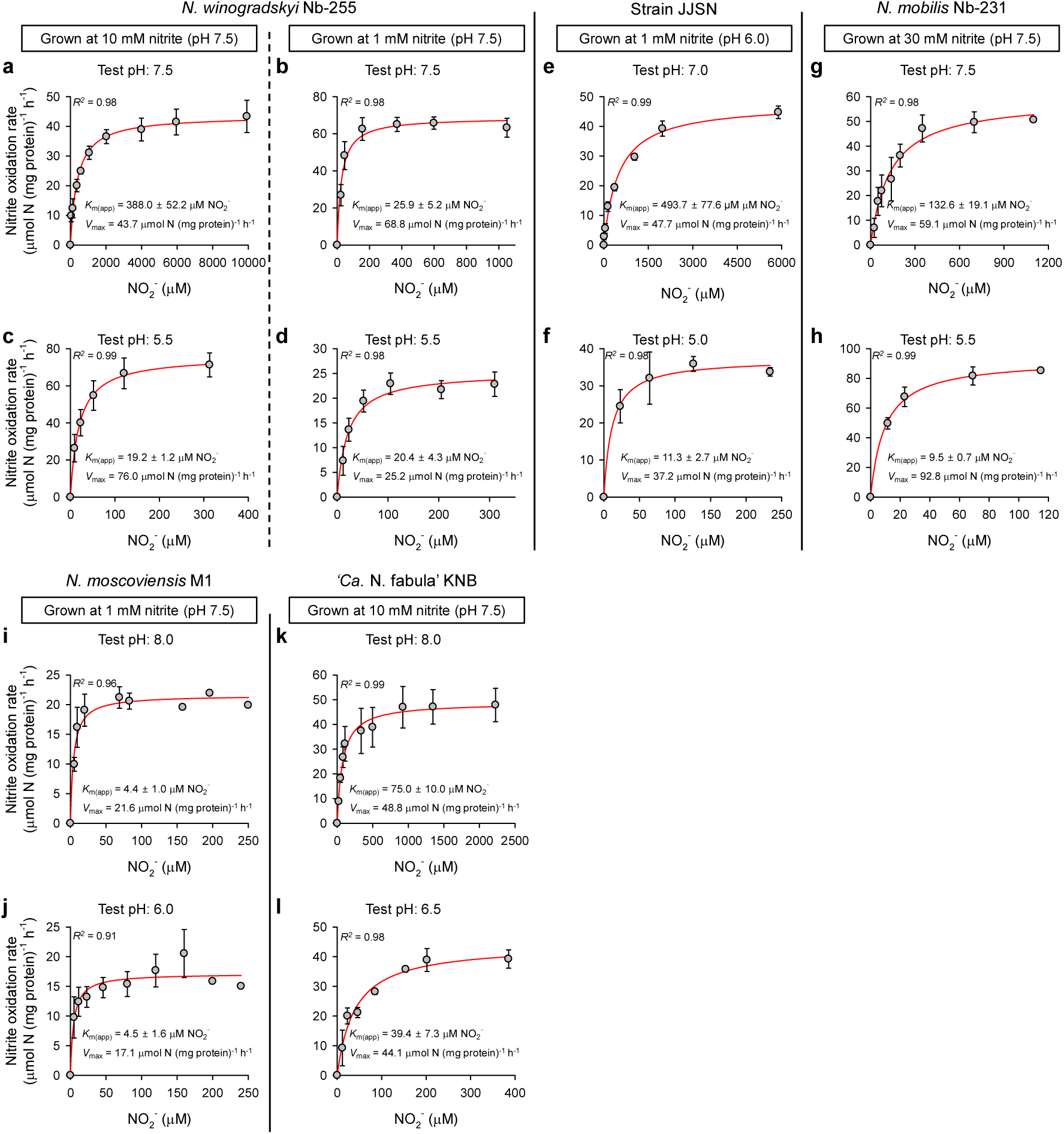
Nitrite oxidation kinetics in NOB strains under different nitrite concentrations and pH. Michaelis-Menten plots for *N. winogradskyi* Nb-255 (**a**–**d**), strain JJSN (**e**, **f**), *N. mobilis* Nb-231 (**g**, **h**), *N. moscoviensis* M1 (**i, j**), and “*Ca*. N. fabula” KNB (**k, l**). Experiments were conducted using mid-log-phase cells exposed to the conditions indicated in each panel. Apparent half-saturation constants (*K*_m(app)_) and maximum nitrite oxidation rates (*V*_max_) were determined by fitting the data to the Michaelis-Menten kinetic equation. The red line indicates the best fit of the data. Standard deviations of the parameter estimates were derived from non-linear regression. Microrespirometry conditions for each strain are provided in **Supplementary Table 3**. Data from all replicate measurements are shown in **Supplementary** Fig. 5.

The mean apparent half-saturation constant (*K*_m(app)_; a measure of affinity) for nitrite oxidation and the maximum oxidation rate (*V*_max_; μmol N mg protein⁻^1^ h⁻^1^) of cells grown with 10 mM nitrite were 388.0 ± 52.2 μM (*n* = 3) and 43.7 ± 1.3 μmol N mg protein⁻^1^ h⁻^1^ (*n* = 3), respectively (Fig. 3a; Supplementary Fig. 5a). These affinity values are consistent with previous reported values for *N. winogradskyi* Nb-255 (*K*_m(app)_ of ∼300 μM) measured under similar conditions (pH 7.5; 9 mM nitrite)^36^. In contrast, when cells grown with 1 mM nitrite surprisingly exhibited a markedly lower *K*_m(app)_ of 25.9 ± 5.2 μM (*n* = 3), but had a *V*_max_ of 68.8 ± 2.5 μmol N mg protein⁻^1^ h⁻^1^ (*n* = 3), which was not significantly different from that of cells grown at higher nitrite concentration (Fig. 3b; Supplementary Fig. 5b). This notably lower *K*_m(app)_ value is comparable to those observed in oligotrophic pNXR-containing NOB such as *Nitrospira* strains^36,40,41^. These findings suggest that cNXR-containing *Nitrobacter* exhibit a higher affinity for nitrite when grown at lower nitrite concentrations.

#### Effect of Test pH on Affinity

To investigate potential pH-dependent changes in nitrite oxidation affinity, independent of growth pH adaptation, we compared the kinetics of *N. winogradskyi* Nb-255 cells cultivated at pH 7.5 under testing conditions of pH 7.5 and pH 5.5. When cells grown with 10 mM nitrite at pH 7.5 were analyzed at the same pH (7.5), a *K*_m(app)_ of 388.0 ± 52.2 μM (*n* = 3) was observed (Fig. 3a; Supplementary Fig. 5a). However, when tested at pH 5.5, the *K*_m(app)_ values significantly decreased to 19.2 ± 1.2 μM (*n* = 6) (Figs. 3c; Supplementary Figs. 5c). This notable shift suggest that cells grown at pH 7.5 can exhibit increased nitrite affinity when exposed to lower pH (pH 5.5) even without prior adaptation to acidic conditions. In the same way, cells cultured with 1 mM nitrite at pH 7.5 showed a *K*_m(app)_ of 20.4 ± 4.3 μM (*n* = 5) when tested at pH 5.5 (Fig. 3d; Supplementary Fig. 5d), which were not significantly different from the *K*_m(app)_ value obtained at test pH 7.5 (*K*_m(app)_ = 25.9 ± 5.2 μM; Fig. 3b; Supplementary Fig. 5b). Comparable pH-dependent increases in nitrite affinity were observed in other cNXR-containing NOB (strain JJSN and *N. mobilis* Nb-231). Both strains exhibited significantly lower *K*_m(app)_ values when cells grown under circumneutral conditions were tested under acidic pH (Fig. 3e–h; Supplementary Figs. 5e–h). In contrast, pNXR-containing strains, *N. moscoviensis* M1 and ‘*Ca.* N. fabula’ KNB showed minimal variation in *K*_m(app)_ values across acidic and neutral pH ranges (Fig. 3i–l; Supplementary Figs. 5i–l). For *‘Ca.* N. fabula’ KNB specifically, the *K*_m(app)_ at pH 6.5 was slightly lower than that at pH 8.0, indicating only moderate pH sensitivity. This limited responsiveness may stem from its distinct NXR variant, which is proposed to form a soluble periplasmic holoenzyme belonging to a sister clade of canonical pNXR^42^.

Across all the tested *Nitrobacter* strains, the *V*_max_ values ranged from 25.2 to 76.0 μmol N mg protein⁻^1^ h⁻^1^, showing minimal variation depending on the initial nitrite concentrations used during cultivation or the tested pH levels. These values were comparable to those reported for pNXR NOB *Nitrospira* strains (18 to 48 μmol N mg protein⁻^1^ h⁻^1^)^36,40,41^. Collectively, these results suggest that *Nitrobacter* strains cultivated under circumneutral conditions exhibit an enhanced affinity for nitrite when exposed to acidic conditions. This trait may be shared among cNXR NOB and could explain the widespread abundance of *Nitrobacter* strains in acidic and nitrite-depleted soil environments^31,37,43^.

#### Comparison of Specific Substrate Affinity

The quantity and kinetic characteristics of the transporter systems strongly influence specific affinity (*a*°)^44^. Thus, for cNXR NOB, *a*° serves as a more ecologically relevant metric than *K*_m_ alone, as it integrates both affinity and maximum uptake capacity, particularly under low-substrate conditions^44^. In this study, *K*_m(app)_ and *a*° (l g cell⁻^1^ h⁻^1^) were compared among isolated or highly enriched NOB possessing either cNXR or pNXR, using data from both this study and previous literature (Fig. 4; Supplementary Table 3)^19,33,36,40,41,45–49^. For *N. winogradskyi* Nb-255, cells cultivated at an initial nitrite concentration of 1 mM showed slightly higher *a*° at pH 7.5 compared to pH 5.5. However, for all other conditions tested in this study, cNXR NOB consistently exhibited higher *a*° values under acidic pH relative to neutral pH (Fig. 4b; Supplementary Table 3). In contrast, as expected, the *a*° values of *N. moscoviensis* M1 and ‘*Ca*. Nitrotoga fabula’ KNB were not remarkably affected by changes in pH (Fig. 4b, Supplementary Table 3), indicating that the substrate affinity of pNXR NOB remains relatively stable across different environmental gradients.

**Fig. 4:**
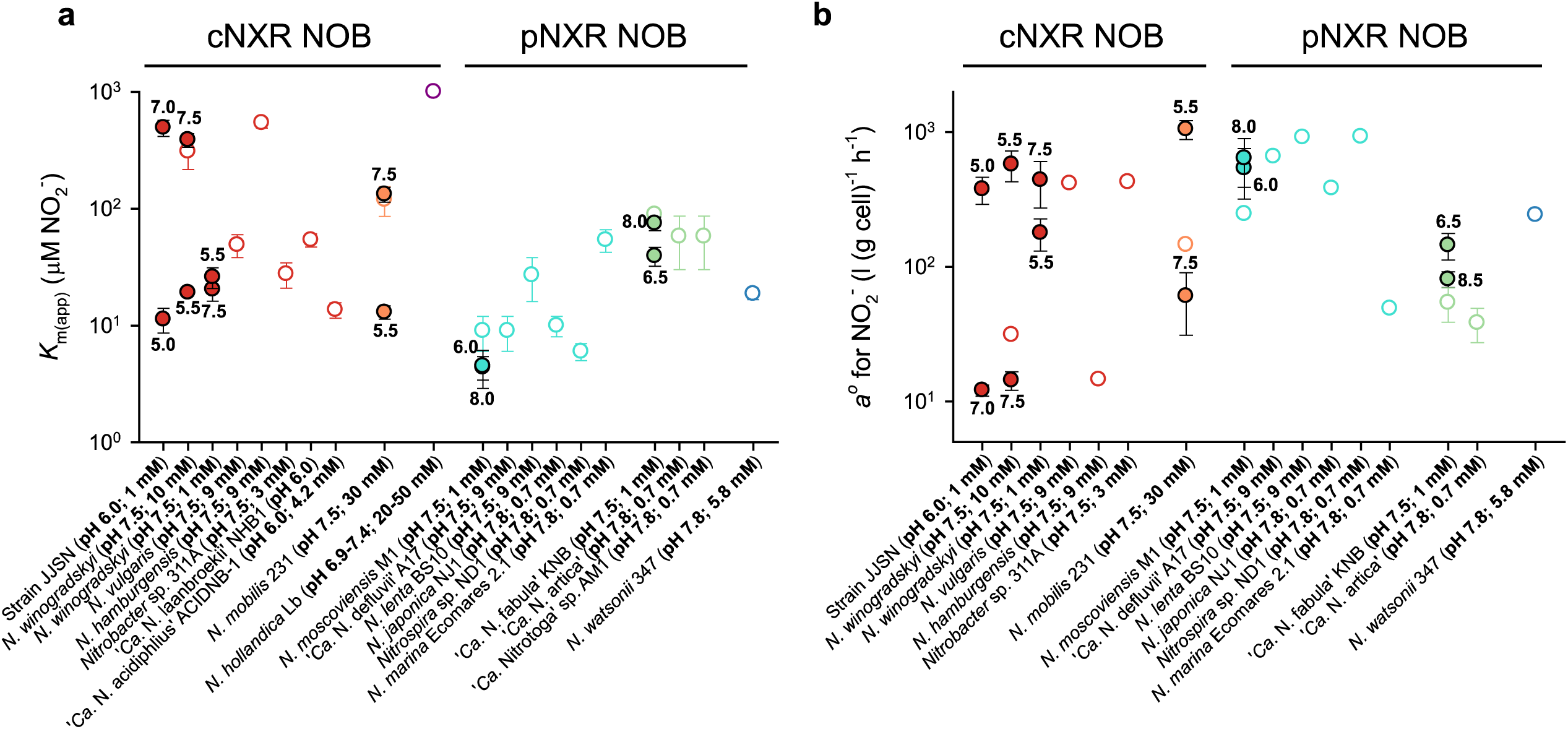
Apparent half-saturation constants (*K*_m(app)_) and specific affinity (*a*°) of NOB strains at different nitrite concentrations and pH. **a**, The *K*_m(app)_ for nitrite. **b**, The *a*° for nitrite. The colours indicate six different genera of NOB strains (*Nitrobacter*: red; *Nitrococcus*: orange; *Nitrolancea*: purple; *Nitrospira*: turquoise; ‘*Ca*. Nitrotoga’: light green; and *Nitrospina*: blue). Filled circles indicate data from this study, while hollow circles represent data from previous studies^19,33,36,40,41,45–49^. The pH of each experiment is shown below each data point. The x-axis denotes strain names and corresponding culture conditions. Error bars represent mean ± s.d. of replicate measurements. The number of replicates (*n*) and corresponding numerical values are provided in **Supplementary Table 3**.

Recently, it was revealed that acid-tolerant *Nitrobacter* strains, ‘*Ca*. N. laanbroekii’ and ‘*Ca*. N. acidophilus’ ACIDNB-1 exhibited relatively higher substrate affinities (*K*_m(app)_ = 54 and 13.6 μM, respectively) when tested at pH 6.0^33,48^. Additionally, *Nitrobacter* strains (*N. vulgaris* and marine *Nitrobacter* sp. Nb-311A) with a relatively high affinity for nitrite (Fig. 4; Supplementary Table 3) have also been reported at circumneutral pH^36,40^. Taken together, our study highlights the need for systematic investigation of the variability of kinetic parameters in cNXR NOB under different environmental conditions, including pH and nitrite concentrations.

The ability of a single microbial species to modulate its substrate affinity in response to environmental conditions is increasingly recognized as a common adaptive strategy among various microbes, challenging the traditional distinction between *r*- and *K*-strategists^44^. For example, *Escherichia coli* cultured in glucose-limited chemostats upregulates high-affinity glucose transporters as nutrient levels decline^50^. Furthermore, it alternates between a high-affinity phosphate transport system (Pst) and a low-affinity system (Pit) based on phosphate availability, thereby optimizing uptake efficiency during periods of starvation^51^. Interestingly, we also found that substrate affinity can increase when cells are exposed to acidic pH, even without prior adaptation to such conditions.

### Nitrite and Nitrate Transporters

The quantity and kinetic properties of substrate transporters are expected to play a significant role in influencing variations in whole-cell nitrite oxidation kinetics in cNXR NOB. Based on genomic analysis, two types of transporters, the Nitrite/Nitrate Porter (NNP; NarK)^52,53^ and the Formate/Nitrite Transport (FNT; NirC) family transporter^54–57^ which are suggested to be responsible for transporting nitrite into the cytoplasm, are conserved across all cNXR NOB (Supplementary Table 4). Alignment of the amino acid sequences of NarK and NirC from *Nitrobacter* revealed conserved residues in the binding site for nitrate and nitrite, as well as within the transport channel regions of these transporter proteins (Supplementary Figs. 6, 7).

NarK is primarily recognized as a nitrite/nitrate antiporter^52^, however alternative mechanisms, such as proton/nitrate symport, have also been proposed^58–61^. Phylogenetic analysis of NarK homologs in cNXR NOB revealed two distinct clades: (i) NarK-c (canonical), affiliated with canonical NarK1 (proton/nitrate symport) and NarK2 (nitrate/nitrite antiport) of heterotrophic bacteria^58,60,61^, and (ii) NarK-n (*Nitrobacter*-type), which is unique to *Nitrobacter* species (Supplementary Fig. 8). Although the transport mechanism of NarK-n has yet to be experimentally confirmed, the conserved genomic linkage of a NarK-n with NXR cluster genes (Supplementary Fig. 9) suggests that it may function similarly to NarK1 and/or NarK2. As for NirC, it is known to function as a nitrite-specific uniporter^57^, typically associated with the nitrite reductase genes (*nirBD*) for the assimilation of nitrite^57,61,62^. However, *nirC* homologs in various NOB genera are not linked to the canonical *nirBD* (Supplementary Fig. 10), suggesting that they may play a role in both nitrite oxidation and nitrite assimilation.

To characterize substrate transporter systems, MR-based whole-cell kinetics analyses were performed using *N. winogradskyi* Nb-255 cells. Excess external nitrate was introduced as a potential inhibitor for nitrate export by the NarK antiporter^63^, which could also interfere with NarK-mediated nitrite uptake^52^. We selected 3 and 10 mM nitrate for 60% inhibition of 1- and 10-mM nitrite-grown cells at pH 7.5, respectively, compared with a no-nitrate control (Supplementary Fig. 11). Notably cells grown with 10 mM nitrite at pH 7.5 showed a remarkable reduction in *V*_max_ compared to *K*_m(app)_ in the presence of 10 mM nitrate at pH 7.5 (Fig. 5a). In the NarK antiport system, excess external nitrate can inhibit nitrite uptake through non-competitive inhibition kinetics. In fact, in bimolecular reactions (A + B → P + Q) similar to antiport systems, elevated concentrations of product P or Q can induce kinetic behavior resembling non-competitive inhibition of substrate A or B^63^.

**Fig. 5:**
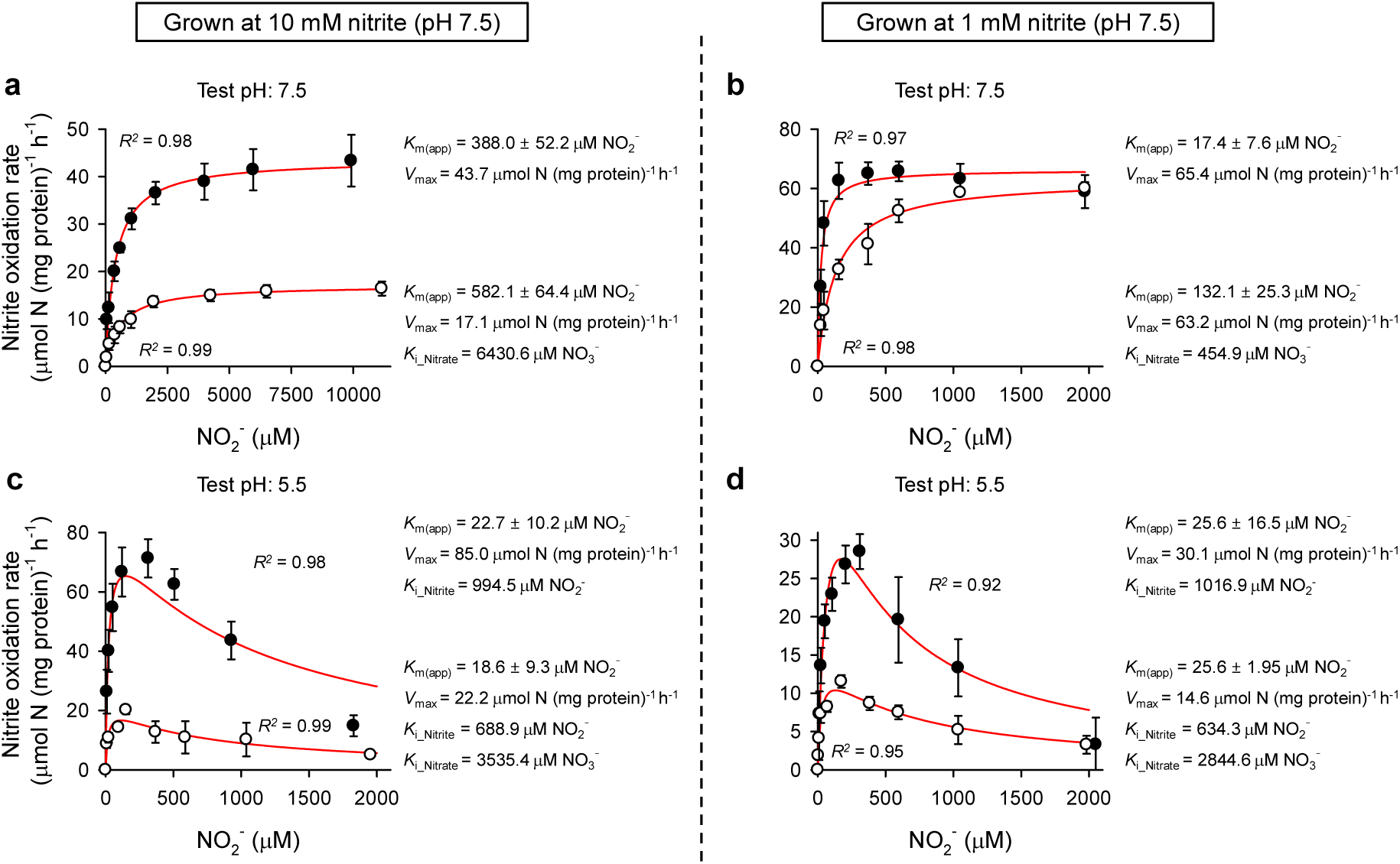
Nitrite oxidation kinetics of *N. winogradskyi* Nb-255 under nitrate inhibition. The nitrite oxidation kinetics of *N. winogradskyi* Nb-255 were analyzed in the presence of nitrate. **a,** Cells grown at 10 mM nitrite (pH 7.5) and tested at pH 7.5; **b,** Cells grown at 1 mM nitrite (pH 7.5) and tested at pH 7.5; **c,** Cells grown at 10 mM nitrite (pH 7.5) and tested at pH 5.5; and **d,** Cells grown at 1 mM nitrite (pH 7.5) and tested at pH 5.5. Cells grown at 10 mM and 1 mM nitrite were exposed to 10 mM and 3 mM nitrate, respectively. In all panels, filled circles represent rates measured in the absence of nitrate. Hollow circles represent rates measured in the presence of nitrate. At pH 5.5 (**c**, **d**), *K*_m(app)_ and *V*_max_ values were derived by fitting data to a substrate inhibition model combined with non-competitive nitrate inhibition, which accounts for nitrite toxicity at high concentrations.

Interestingly, *N. winogradskyi* Nb-255 cells grown with 1 mM nitrite (pH 7.5) demonstrated competitive inhibition-like kinetics in the presence of 3 mM nitrate at the same pH. This behavior can be attributed to the slow recovery of the nitrite oxidation rate upon adding more nitrite (Fig. 5b). One possible explanation for this phenomenon could be a decoupling of nitrite and nitrate transporters. Under this condition, the NirC transporter may mediate nitrite uptake in cells cultivated with 1 mM nitrite. *E. coli* possessing NirC can take up nitrite ∼10× faster under nitrite-limited conditions compared to strains that lack NirC, highlighting its significance when nitrite is limited^58^. Since NirC’s pore binds and translocates both nitrate and nitrite, these anions compete for access to the same transport pathway^57^, making nitrate act as a competitive inhibitor to nitrite. Notably, NirC generally exhibits a higher substrate affinity^54–56^ than NarK^61,64^, this may explain the higher nitrite affinity observed in *N. winogradskyi* Nb-255 cells grown at 1 mM nitrite compared to those grown at 10 mM nitrite (Fig. 3a, b; Fig. 5a, b). We propose that under this condition, NarK may play a role in nitrate efflux as a proton/nitrate symporter, which could also contribute to proton motive force and pH homeostasis^65^.

When cells grown with 1 and 10 mM at pH 7.5 were exposed to an acidic condition (pH 5.5) in the absence of nitrate, they showed strong inhibition of nitrite oxidation at higher nitrite concentration (Fig. 5c, d), reflecting typical substrate inhibition kinetics, which may be due to HNO_2_ toxicity. At pH 5.5, cells exhibited increased nitrite affinity (lower *K*_m(app)_), which occurred without prior adaptation to acidic conditions and remained independent of nitrate inhibition (Fig. 3; Fig. 5c, d). This observation suggests a combination of the following mechanisms: enhanced NarK antiport activity under acidic condition^51^ and increased passive diffusion of HNO_266_. Increased antiport activity of NarK at acidic conditions was observed in NarK-reconstituted proteoliposomes^51^. Acid-driven HNO_2_ diffusion into the cytoplasm may contribute to intracellular nitrite accumulation at a circumneutral cytoplasmic pH, which could serve as a sink for HNO_2_. While the involvement of high-affinity NirC at acidic pH is conceiveable^58^, competitive inhibition of nitrate at low nitrite concentrations may be obscured in kinetic analyses (Fig. 5c, d) due to increased activity of NarK and enhanced diffusion of HNO_2_ under these conditions. Conversely, at higher nitrite concentrations, the nitrite oxidation rates are similar (Fig. 5c, d), as indicated by competitive inhibition (see Fig. 5b).

In both RT-qPCR and transcriptomic analysis (Supplementary Fig. 12; Supplementary Datasets 2, 3), *narK*-n (Nwi_0779) was highly and constitutively expressed across all tested conditions, and only *nirC* (Nwi_3006) expression was upregulated at lower nitrite concentrations and acidic condition (Supplementary Datasets 2, 3; Supplementary Fig. 12), indicating its principal role as a key high affinity nitrite transporter. Together, these findings imply that modulation of transporter activity, such as through allosteric or covalent modification, is a core factor in dynamic adaptation to environmental changes, as has been well demonstrated in bacteria and plants^67,68^. Regulatory proteins such as nitrogen regulatory protein P-II (Nwi_2931), upregulated at acidic pH (Supplementary Dataset 2), may play a role in modulating the activity of the membrane-bound transporter^67,69,70^. However, the detailed mechanistic studies on transporter activity modulation in cNXR NOB remain unexplored and require further investigation. Overall, these findings propose a schematic model for the dynamic kinetics of nitrite oxidation mediated by transporter systems in *Nitrobacter*, as illustrated in Fig. 6.

**Fig. 6.**
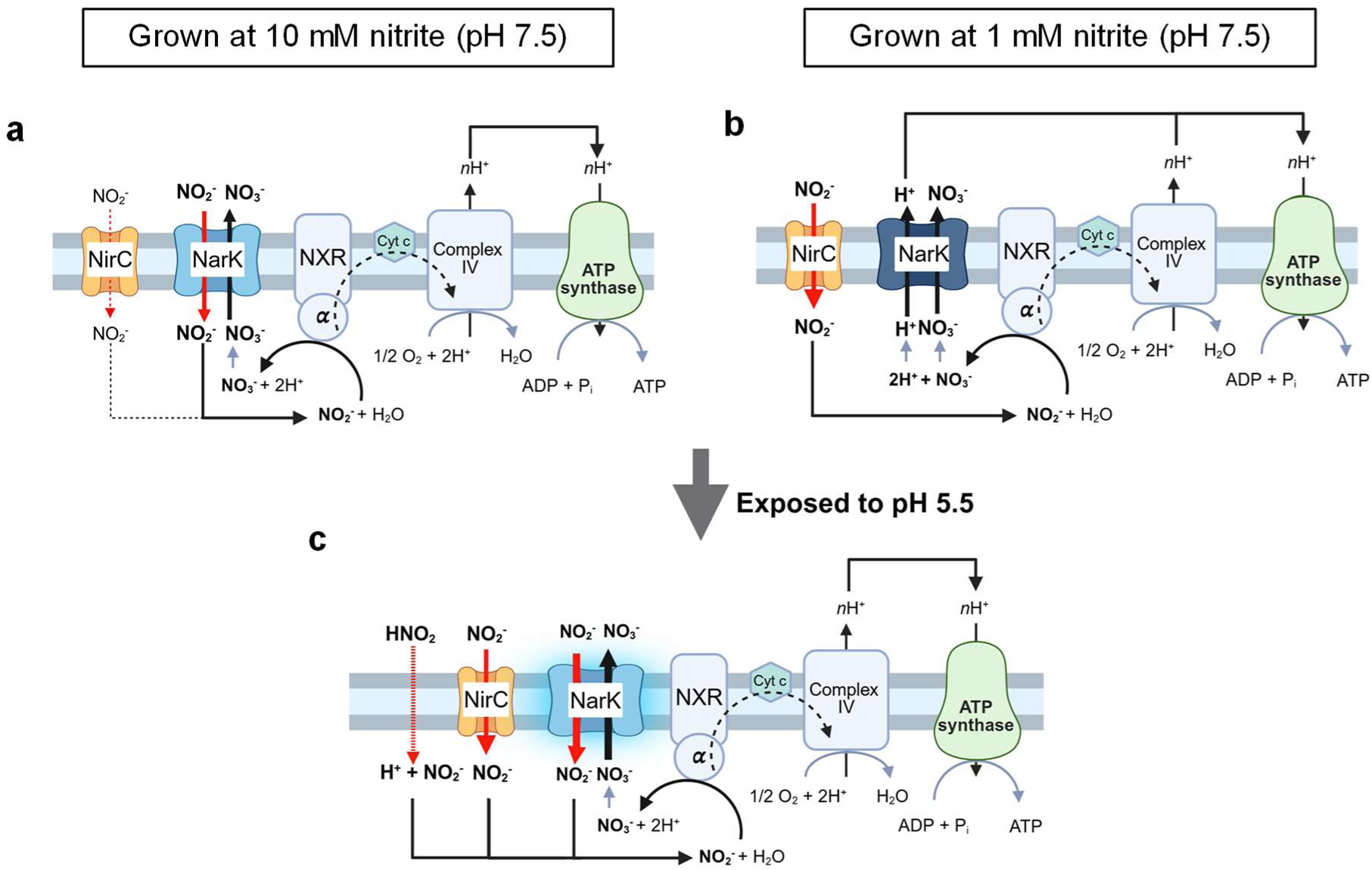
Schematic model of transporter systems influencing nitrite oxidation kinetics in Nitrobacter. **a,** Cells grown at 10 mM nitrite (pH 7.5). Nitrite uptake is primarily mediated by low-affinity NarK, likely functioning as a nitrite/nitrate antiporter. NirC may transiently facilitate intracellular nitrite accumulation, thereby triggering cNXR activity. **b,** Cells grown at 1 mM nitrite (pH 7.5). Under low nitrite conditions, nitrite uptake is predominantly mediated by NirC, which may have a higher affinity for nitrite than NarK^54–56,61,64^. Polytopic NarK may act as a candidate nitrate exporter via H⁺/nitrate symport, potentially contributing to pH homeostasis. **c**. Cells pre-cultured at pH 7.5 (with 10 or 1 mM nitrite) and exposed to pH 5.5. The increased nitrite affinity observed at pH 5.5 as shown in Fig. 3, even without prior acid adaptation, can be explained by the following combined processes: 1) Positive modulation of NarK antiporter activity at acidic pH^52^, 2) Increased passive diffusion of HNO_2_ across the membrane which may cause intracellular nitrite accumulation, and 3) Increased high-affinity import of nitrite by NirC under acidic and low nitrite conditions via upregulated gene expression.

Our study suggests that differential use of transporters, combined with the immediate regulation of pre-existing transporters, is involved in this lifestyle switch, as indicated by the kinetic and transcriptomic analyses. This mechanism may play a pivotal role in maintaining high nitrite oxidation rates in acidic soils, thereby contributing to nitrogen retention by preventing the abiotic decomposition of nitrite in these conditions^34,35^ and minimizing its release into the atmosphere as HONO gas (nitrous acid). Furthermore, this adaptive kinetic plasticity is likely extended to other cNXR NOB in response to varying environmental conditions.

### Conclusions

*Nitrobacter* strains exhibit a shift in optimal pH for nitrite oxidation toward acidic conditions at lower nitrite concentrations. Whole-cell kinetics revealed a higher nitrite affinity in cells grown at lower nitrite concentrations. Furthermore, cells grown at neutral pH also exhibit increased nitrite affinity under acidic test conditions, even without prior adaptation to acidity. This adaptive plasticity is likely mediated by the regulations of gene expression levels and the kinetic behavior of nitrite transporters, such as NirC and NarK, which are conserved and universally present across all known cNXR NOB. Understanding the regulation of transporter activity that governs such kinetic responses is crucial for unraveling the adaptive mechanisms of cNXR NOB. Notably, ammonia oxidation can locally reduce pH in weakly buffered soil microniches, favoring NOB with high nitrite affinity under acidic conditions. *Nitrobacter* coexisting with AOM may adapt to such microacidic niches, highlighting the role of kinetic plasticity in response to spatial pH heterogeneity. Therefore, this study provides valuable insights into the niche specialization of cNXR NOB, shedding light on the complex spatial distribution patterns and temporal population dynamics of coexisting NOB and their interactions with other nitrogen cycle microbes, particularly AOM, across diverse environments.

Although the total nitrite (NO_2_⁻) ratio is still high (HNO_2_ < 1%) at acidic conditions (e.g., pH 5.0), ammonia oxidation is constrained by the steep decline in bioavailable ammonia (NH_3_), resulting in markedly lower overall substrate (nitrite) concentration for NOB at low pH. Under such conditions, *Nitrobacter* exhibits enhanced nitrite affinity that is comparable to that of oligotrophic pNXR NOB such as *Nitrospira*, providing a mechanistic explanation for its persistence in nitrite-depleted, acidic soils. Furthermore, cytoplasmic localization of NXR may confer protection against environmental stressors, such as extreme pH, HNO_2_, and heavy metals. Thus, despite being traditionally viewed as copiotrophs, cNXR NOB exhibit physiological flexibility that enables niche expansion across diverse environments.

However, *Nitrospira* often dominates numerically, possibly due to additional advantages such as scalar proton generation via pNXR and the use of the more energy-efficient rTCA cycle over the Calvin-Benson-Bassham cycle in *Nitrobacter*. These combined traits suggest that nitrite affinity alone does not determine ecological success; instead, multiple metabolic and energetic factors contribute to shaping niche partitioning among NOB taxa. Our study has broad implications for predicting NOB responses to fluctuating environmental conditions and for incorporating their contributions into nitrogen cycle models and ecosystem nutrient budgets.

### Newly isolated Nitrobacter species

The isolated strain JJSN is a novel species of the genus *Nitrobacter* within the order *Hyphomicrobiales*, and we propose the following candidate status:

#### Taxonomy. (i) Etymology

The taxonomy for ‘*Candidatus* Nitrobacter acidaffinis’ sp. nov. is as follows: L. adj. acidus, acidic; L. adj. affinis, related to, associated with; N.L. adj. acidaffinis, acid-associated (referring to acid-tolerant nature of the strain).

**(ii) Locality.** Isolated from an acidic forest soil (pH 4.6) in Jeju Island, Republic of Korea.
**(iii) Diagnosis.** The strain oxidizes nitrite efficiently at pH as low as 5.0, exhibits cytoplasmic NXR localization, and harbors a distinct *narK* gene within the NXR cluster. The strain exhibits a dynamic shift in nitrite oxidation affinity, depending on substrate concentration and pH, demonstrating its capacity for physiological plasticity. Phylogenomic analysis based on 71 core genes and ANI values (< 90% to known *Nitrobacter* species) supports its designation as a novel species within the genus *Nitrobacter*.

## Materials and Methods

### Enrichment and isolation of acid-adapted NOB

To cultivate the acid-adapted NOB strains, soil samples were collected from a depth of 30 cm in forest soil located in Jeju, Korea (33°25’6.85”N, 126°39’36”E) in January 2022. Soil pH rnaged from 4.2 to 5.0 (*n* = 3) using a 1:5 (w/w) soil-to-water ratio (Supplementary Table 1). Nitrite was below the detection limit, while ammonia and nitrate concentrations were in the micromolar and millimolar ranges, respectively, in acidic soils. (Supplementary Table 1). Enrichment culture was initiated by inoculating 0.5 g of the forest soil sample into 100 mL of an autotrophic NOB medium (as described in Nowka *et al*.^36^), adjusted to pH 5.0 and incubated at 25 °C. Cultures were supplied with 100 µM NaNO_2_ to minimize the toxicity of HNO_2_, under acidic pH conditions^38^. Nitrite and nitrate concentrations were determined colorimetrically using the Griess assay in 96-well flat-bottom microplates, as previously described^71^. For nitrite measurement, 50 μL of sulphanilamide solution (5 g L⁻^1^ in 2.4 M HCl) was added to 50 μL of the sample or standard, followed by the addition of 50 μL of naphthylethylenediamine solution (3 g L⁻^1^ in 0.12 M HCl). Nitrate concentrations were measured using a nitrate reduction method by adding 15 μL of vanadium (III) chloride solution (10 g L⁻^1^ in 1 M HCl). Standards were prepared in duplicate, using NaNO_x_, and ranged from 5.0 to 500 µM. Absorbance was recorded at a wavelength of 540 nm using a Spectra Max M2 plate reader (Molecular Devices, CA, USA). The increase in nitrite oxidation activity was accompanied by a corresponding increase in the copy number of the 16S rRNA gene copy number, as assessed by quantitative PCR (qPCR) of extracted DNA. qPCR was performed under the following conditions: initial denaturation at 95 °C for 3 min; 40 cycles of 95 °C for 30 s, 55 °C for 30 s, and 72 °C for 30 s, with a final extension at 72 °C for 5 min. Amplification efficiency was 94% for target gene, and qPCR *r*^2^ calibration values were greater than 0.99.

The enrichment culture was regularly transferred at 1% (v/v) into fresh medium containing 100 µM NaNO_2_ approximately once per week, and a pure culture designated strain JJSN was obtained through subsequent endpoint serial dilution in deep-well 96-well plates. All cultures were maintained in the dark, without shaking, at 25 °C. The absence of heterotrophic contaminants was confirmed by bacterial 16S rRNA gene-based amplicon sequencing^72^. In addition, absence of heterotrophic growth on nutrient-rich solid agar plates (R2A broth and tryptic soy broth) was confirmed weekly after one week of incubation.

### Cultivation of previously isolated NOB strains

This study additionally utilized four distinct NOB isolates from previous studies, each representing a different genus: *Nitrobacter winogradskyi* Nb-255, *Nitrococcus mobilis* Nb-231, *Nitrospira moscoviensis* M1, and ‘*Ca*. Nitrotoga fabula’ KNB (Supplementary Table 5), along with the acid-adapted nitrite-oxidizing bacterial strain newly isolated in this study (strain JJSN). All strains were cultivated in batch culture under the conditions listed in Supplementary Table 5. Briefly, cells were grown in 25 cm³ cell culture flasks (SPL Life Sciences, Republic of Korea) at their optimal temperature, in the dark, without shaking, unless otherwise specified. Substrate was supplied as NaNO_2_ from sterilized stock solutions. To maintain culture pH, 2 mM sterile NaHCO_3_ and 3 mM MES or HEPES buffer were added, titrated with NaOH as necessary.

The pH, temperature, and nitrite concentrations were selected based on the physiological characteristics of each strain (see Supplementary Table 5). Considering abiotic decomposition of HNO_2_ below pH 4.5 (see Supplementary Fig. 2), pH levels from 5.0 to 9.0 (in 0.5-unit intervals) were tested for the NOB strains with 10–10,000 µM nitrite. Due to the acid adaptation of strain JJSN, pH 4.5 was used for its nitrite oxidation activity (see Supplementary Fig. 2). Each experiment was performed in biological triplicate using a 1% (v/v) inoculum from nitrite-grown cultures, with daily sampling. The nitrite oxidation rate was calculated using the following equation (2):

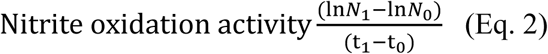

where (ln*N*_1_ − ln*N*_0_) represents the change in the natural logarithm of nitrite concentration over the time interval (t_1_ − t_0_). At least three time points were used to calculate the nitrite oxidation rate.

### Kinetic analysis

Cellular substrate oxidation kinetics were determined from instantaneous, substrate-dependent oxygen uptake measurements as previously described. Briefly, measurements were performed using a microrespirometry (MR) system, equipped with a PA 2000 picoammeter and a 500 µm tip-diameter OX-MR oxygen microsensor (Unisense, Denmark), which was continuously polarized for at least 24 hours before use. Active NOB cells were harvested (5,000 × *g*, 20 min, 25 °C) from nitrite-replete (mid**-**log**-**phase) active cultures, using 30 kDa-cutoff Amicon Ultra-15 centrifugal filter units (Merck Millipore, Germany). Concentrated cells were washed two to three times and resuspended in a substrate-free medium prior to MR experiments.

The MR experiments were conducted at the optimal growth temperature with varying pH conditions (Supplementary Table 3). For all MR experiments, glass MR chambers containing glass-coated magnetic stir bars were filled headspace-free, sealed with MR injection lids, and submerged in a recirculating water bath, under constant stirring (200 rpm). An OX-MR microsensor was inserted into each MR chamber and left to equilibrate (for approximately 1 h). Exact temperatures and pH conditions used for each strain and experiment are provided in Supplementary Table 3. Stable background sensor signal drift was measured for at least 30 min before initial substrate injections, and the background oxygen consumption rate was subtracted from the measured oxygen uptake rates. NaNO_2_ stock solutions were prepared in non-inoculated medium adjusted to the same composition as the cultivation medium and to the pH of the MR testing conditions, and were injected into the MR chambers using Hamilton syringes (25 µL; Hamilton, USA). Both single- and multiple-trace oxygen uptake measurements were performed. For single-trace measurements, a single substrate injection was performed, and oxygen uptake was recorded until the substrate was depleted. For multiple-trace measurements, multiple injections of varying substrate concentrations were performed. Once stable, discrete slopes of oxygen uptake were calculated following each substrate injection. Immediately following oxygen uptake zmeasurements, the nitrite and nitrate concentrations and pH of the MR chamber contents were determined.

For nitrate inhibition kinetics, 1 mM- and 10 mM-nitrite-grown *N. winogradskyi* Nb-255 cells were used in each condition (Supplementary Table 3). First, inhibitory nitrate concentrations that caused approximately 60% inhibition of nitrite oxidation relative to no-nitrate control were determined. For this, after incubating for approximately 1 h in substrate-free medium at each pH (5.5 and 7.5), 375 µM NaNO_2_ was injected into the MR chamber. Once a stable slope of oxygen uptake was obtained, multiple injections of increasing NaNO_3_ concentrations (1, 3, and 10 mM) were performed in the same MR chamber. Then, for the nitrate inhibition kinetics, multiple trace measurements were performed in the presence of specific nitrate concentrations, as described above.

For protein determination, the cells used for kinetic analysis were stored at -20 °C. Cells were lysed with the Bacterial Protein Extraction Reagent (Thermo Scientific, USA), and the total protein content was determined photometrically with the Pierce bicinchoninic acid (BCA) Protein Assay Kit (Thermo Scientific, USA) as per the manufacturer’s instructions.

### Calculation of kinetic parameters

*K*_m(app)_ and *V*_max_ were calculated from both single- and multiple-trace substrate-dependent oxygen uptake measurements. Nitrite oxidation rates were calculated from oxygen uptake measurements using a substrate-to-oxygen consumption ratio of 2:1 (Eq. 1). Nitrite oxidation rates were fitted to a Michaelis–Menten model using the equation (Eq. 3):

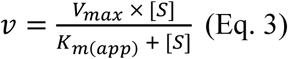

where *v* is the reaction rate (μmol h⁻^1^), *V*_max_ is the maximum reaction rate (µmol N mg protein⁻^1^ h⁻^1^), *S* is the nitrite concentration (µM), and *K*_m(app)_ is the reaction half saturation concentration (µM). A non-linear least-squares regression analysis was performed to estimate *K*_m(app)_ and *V* ^73^ by fitting the experimental data. Under acidic conditions, data were collected at non-inhibitory nitrite levels, unless otherwise noted. The *K*_m(app)_ for nitrite was calculated for each strain at each tested pH condition and nitrite concentration supplied during batch growth. The specific substrate affinity (*a*°; l g cell⁻^1^ h⁻^1^) of each pure culture strain was calculated using the equation:

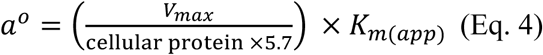

Where the *V*_max_ is normalized to the protein concentration (g l⁻^1^) of the culture in the MR chamber, and the factor of 5.7 g wet cell weight per g of protein was used for all NOB^74^. The *a*° for nitrite was calculated using the respective *K*_m(app)_ for nitrite.

Nitrate inhibition kinetics can be described by the Michaelis–Menten equation incorporating either the non-competitive inhibition model (Eq. 5), the competitive inhibition model (Eq. 6), or both substrate and non-competitive inhibition models (Eq. 7). Inhibitory concentrations (*K*_i_ values) were determined by fitting the three possible Michaelis-Menten inhibition models (Eqs. 5–7) to the experimental data and selecting the model that provided the best fit.

*I* is the nitrate concentration (µM) supplemented, *K*_m(app)_ and *V*_max_ values for nitrite were determined experimentally using non-inhibitory nitrite concentrations at an acidic pH of 5.5.

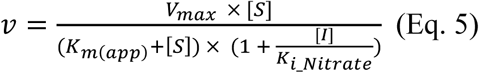

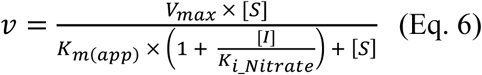

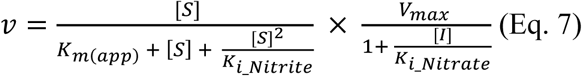

### Genomic and phylogenetic analysis

Biomass for DNA extraction from strain JJSN was harvested from 1,000 mL aliquots of the cultures by filtration, using mixed cellulose ester (MCE) filters (Advantec Mfs Inc., Dublin, CA, USA) with a diameter of 47 mm and a pore size of 0.2 µm. The collected filters were ground in liquid nitrogen using a mortar and pestle. Genomic DNA was extracted using a modified hexadecyltrimethylammonium bromide (CTAB)-based protocol^75^. Long-read sequencing libraries were prepared with the Native Barcoding Kit 24 V14 (SQK-NBD114.24). The genome of strain JJSN was sequenced using the MinION R10.4.1 flow cell (FLO-MIN114, Oxford Nanopore Technologies) at laboratory and the Illumina NovaSeq6000 (2 × 150 bp) at LabGenomics (Seongnam, Republic of Korea). Base calling was carried out with Dorado (v0.8.0), integrated within the MinKNOW software (v23.04.6). A total of 662,835,824 bp were generated from 1,067,406 reads (maximum read length: 57,482 bp), with a median quality score of 16.02, corresponding to an accuracy of approximately 97.5%. De novo assembly of MinION long reads was performed using Flye (v2.9.2)^76^, and the resulting consensus contigs were polished with Illumina short reads using Polypolish (v0.5.0)^77^ and POLCA (v4.0.5)^78^ to improve base-level accuracy. Genome circularity was confirmed during the Trycycler pipeline assembly and further validated by mapping Illumina reads in a reverse orientation.

Pylogenomic analyses were conducted based on genomes available in the NCBI database. A total of 29 genomes and metagenome-aseembled genomes (MAGs) were included in a concatenated marker gene analysis: 24 previously published genomes from Nitrobacter, the genome of strain JJSN (this study), and 4 genomes from the family Nitrobacteraceae (*Bradyrhizobium, Rhodopseudomonas, Afipia*) used as outgroup taxa. Seventy-one bacterial single-copy marker genes were extracted from the genomes using HMMER v3.3.2^79^ with the HMM profiles bundled in anvi’o v7.1. The extracted gene sequences were aligned with MUSCLE v3.8.1551^80^ and concatenated using anvi’o v7.1. A maximum likelihood phylogenetic tree was constructed using IQ-TREE (v1.6.12)^81^. The LG+F+R4 model was selected using ModelFinder Plus^82^. Trees were visualized in iTOL v7.1^83^.

For NxrA analyses, NxrA protein sequences (> 1200 amino acids) from *Nitrobacter* were obtained from the NCBI database based on their association with the NXR gene cluster. The NxrA sequence from the genus *Nitrococcus* was used as the outgroup. A maximum likelihood phylogenetic tree was constructed using IQ-TREE (v1.6.12)^81^ with the K3Pu+F+I+R3 substitution model.

In parallel, genome-wide similarity analyses were conducted to assess taxonomic boundaries based on ANI and AAI, calculated as described in previous studies^84,85^. Genomes from members of the *Nitrobacteraceae* family (*Nitrobacter*, *Bradyrhizobium*, *Rhodopseudomonas*, and *Afipia)* were obtained from the NCBI database and filtered using a cutoff of 80% identity to strain JJSN in both ANI and AAI. Pairwise ANI and AAI values were clustered and visualized as heatmaps using R (v4.2.3) with the ggplot2 (v3.5.1) and reshape2 (v1.4.4) packages. Species-level boundaries were determined using a 95% threshold for ANI, as proposed in previous studies^86,87^.

### Nitrite/nitrate transport gene analysis

We analyzed the presence of putative nitrite transporter genes (*nirC*, *narK*, and *nrt*ABC) in 26 publicly available genomes of NOB harboring either cytoplasmic (cNXR) or periplasmic (pNXR) nitrite oxidoreductase. These genomes were derived from isolated strains or highly enriched cultures representing six distinct genera: *Nitrobacter*, *Nitrococcus*, *Nitrolancea*, *Nitrospira*, *Nitrospina*, and ‘*Ca*. Nitrotoga’.

Genes potentially encoding nitrite transporters were identified through domain-based screening using the Conserved Domain Database (v3.18)^88^ and Pfam (v34.0)^89^, implemented via the InterProScan software package (v5.56−89.0). Putative hits were subsequently curated through manual inspection to confirm functional annotations. A phylogenetic tree was reconstructed using IQ-TREE with the LG+F+R4 model. Multiple sequence alignments (MSAs) of *N. winogradskyi* Nb-255 NarK and NirC with homologous sequences were performed using Clustal Omega^90^ with default option and visualized with ESPript 3^91^.

### RNA extraction and gene expression analyses

For both RT-qPCR and transcriptome analyses, five biological replicates of total RNA were extracted from *N. winogradskiy* Nb-255 cells cultured under different nitrite and pH conditions. Initially, 1,000 mL of DSMZ 756c medium^36^ was used to cultivate *N. winogradskiy* Nb-255 under each condition. Total RNA was collected from cells in liquid cultures after approximately 50% of the nitrite had been oxidized. The cells were harvested by filtration, using mixed cellulose ester (MCE) filters (Advantec Mfs Inc., Dublin, CA, USA) with a diameter of 47 mm and a pore size of 0.2 µm. The filters were individually processed by grinding with a mortar and pestle in liquid nitrogen. Total nucleic acids (DNA and RNA) were extracted from each sample using a modified CTAB reagent^75^. The extracted nucleic acids were then processed with the AllPrep DNA/RNA Mini Kit (Qiagen, Germany) to separate RNA from DNA, following the manufacturer’s instructions. RNA purification was performed using the ezDNase kit (Thermo Fisher Scientific, USA), and RNA concentrations were measured using a Qubit 4 fluorometer (Thermo Fisher Scientific, USA). The absence of residual genomic DNA was confirmed via PCR amplification using universal 16S rRNA gene primers for 30 cycles.

For RT-qPCR, total RNA was extracted from cells grown at 1 mM and 10 mM NaNO_2_ at pH 7.5. cDNA synthesis was performed using the SuperScript IV VILO ezDNase kit (Thermo Fisher Scientific, USA). The expression of key nitrite transporter genes (*nirC*, *narK-* n, and *narK-*c) was relatively quantified using *recA* and *nxrA* as housekeeping genes. Primer sequences are listed in Supplementary Table 6. Amplification was carried out under the following conditions: 95 °C for 3 min; 40 cycles of 95 °C for 30 s, 55 °C for 30 s, and 72 °C for 20 s; and a final extension at 72 °C for 5 min. Amplification efficiencies ranged from 80% to 94%, with qPCR calibration *r*² values exceeding 0.99.

For transcriptomic analysis, RNA from cells grown at 10 mM (pH 7.5), 1 mM (pH 7.5), and 1 mM (pH 6.0) NaNO_2_ was submitted to Phyzen (Seongnam, Korea). Libraries were prepared using the Nugen Universal Prokaryotic RNA-Seq Library Preparation Kit without rRNA removal and sequenced with Illumina NovaSeq 6000 platform. Quality was assessed via FastQC (v0.11.8)^92^, trimming with Trimmomatic (v0.36)^93^ using SLIDINGWINDOW:4:15 LEADING:3 TRAILING:3 MINLEN:38 HEADCROP:13, and filtered with SortMeRNA (v2.1)^94^. Reads were aligned to the genome using Bowtie2 (v2.4.4)^95^, and counts obtained with HTSeq (v0.12.3)^96^. Gene expression was normalized as transcripts per kilobase million (TPM) and differential expression assessed using DEseq2 (v1.42.1)^97^ in R (v4.3.1)^98^.

### Statistical analysis

All statistical analyses were conducted using the R statistical software (version 4.2.3), RStudio (version 2023.03.3), and SigmaPlot 10.0 (Systat Software Inc., San Jose, CA, USA). One-way or two-way ANOVA followed by Tukey’s test was used to assess differences among groups under different conditions.

### Data availability

The complete genome sequence of strain JJSN was deposited in the National Center for Biotechnology Information (NCBI) GenBank (PRJNA1297094). The transcriptomic sequence of strain Nb-255 was deposited in the NCBI GenBank (PRJNA1295709).

## Acknowledgments

This work was supported by the NRF (National Research Foundation of Korea) grant funded by the Korean government (Ministry of Science and ICT) (2021R1A2C3004015), the Basic Science Research Program through NRF funded by the Ministry of Education (2020R1A6A1A06046235), and the Korea Institute of Marine Science & Technology Promotion (KIMST) funded by the Ministry of Oceans and Fisheries (RS-2025-02307311). J-H.G. was supported by the NRF grant funded by the Korean government (Ministry of Science and ICT) (RS-2023-00213601). M.-Y.J. was supported by the NRF grant funded by the Korean government (Ministry of Science and ICT) (RS2025000518246). MW, HD, and KK were supported by the Austrian Science Fund (FWF) Cluster of Excellence Microbiomes drive Planetary Health (10.55776/COE7). Dr. Suk-Hyung Ko of the World Heritage Office, Jeju Special Self-Governing Provincial Government, assisted with soil sampling in the forests of Jeju Island.

## Author contributions

U.-J.L., J.-H.G., M.-Y.J., and S.-K.R. designed research. U.-J.L., C.A., S.L., J.-S.Y., O.-J.S., and H.J.C. performed research. U.-J.L., J.-H.G., Z.-X.Q., K.K., H.D., M.W., and S.-K.R.analyzed data. U.-J.L., J.-H.G., M.-Y.J., M.W., and S.-K.R. wrote the manuscript with contributions and comments from all co-authors.

