## Supplementary Information for "Kinetic Plasticity of Nitrite-Oxidizing Bacteria Containing Cytoplasmic Nitrite Oxidoreductase"

[1]Department of Biological Sciences and Biotechnology, Chungbuk National University, Cheongju, Republic of Korea. [2]School of Life Sciences, Fudan University, Shanghai, China. [3]Centre for Microbiology and Environmental Systems Science, University of Vienna, Vienna, Austria. [4]The Comammox Research Platform, University of Vienna, Vienna, Austria. [5]Center for Microbial Communities, Department of Chemistry and Bioscience, Aalborg University, Aalborg, Denmark. [6]Department of Science Education, Jeju National University, Jeju, Republic of Korea. [7]Interdisciplinary Graduate Program in Advance Convergence Technology and Science, Jeju National University, Jeju, Republic of Korea

**This document includes:**

Supplementary Tables 1 to 6

Supplementary Figures 1 to 12

Legends for Supplementary Datasets S1 to S3

Supplementary References

### Supplementary Table 1–6

**Supplementary Table 1. Physicochemical properties of Jeju Forest soil used for the isolation of strain JJSN.**

| Property | Value |
|---|---|
| Coordinates | 33° 25' 6. 85" N<br>126° 39' 36" E |
| Soil texture | Sandy loam |
| Water content (%) | 36.6 ± 4.3 |
| pH [1:5] | 4.6 ± 0.4 |
| Electrical conductivity (μs $cm^{-1}$) | 33.5 ± 6.5 |
| Organic Matter (g $kg^{-1}$) | 58.2 ± 4.8 |
| Total phosphate (g $kg^{-1}$) | 1.7 ± 0.5 |
| Total Nitrogen (g $kg^{-1}$) | 1.6 ± 0.2 |
| $NH_4^+$-N (mg $kg^{-1}$) | 2.8 ± 0.1 |
| $NO_2^-$-N (mg $kg^{-1}$) | N.D. (< 0.01)* |
| $NO_3^-$-N (mg $kg^{-1}$) | 41.3 ± 0.3 |
| Potassium ($K^+$) (cmol(+) $kg^{-1}$) | 4.3 ± 1.5 |

*N.D., not detected; detection limit = 0.01 mg $kg^{-1}$

**Supplementary Table 2. Genomic features of strain JJSN**

| Attribute | Value |
|---|---|
| Environment | Forest soil |
| Genome size (Mb) | 4.12 |
| Coding DNA sequences | 3929 |
| Average CDS length (bp) | 874 |
| Number of scaffolds | 2 |
| Average G+C content (%) | 60.95 |
| Number of rRNAs | 3 |
| Number of tRNAs | 53 |
| *nxrA* gene | 3 |
| *nxrB* gene | 2 |

**Supplementary Table 3. Kinetic constants of nitrite oxidation in NOB, including apparent half-saturation constants ($K_{m(app)}$), maximum specific activity ($V_{max}$), and specific affinity ($a^o$).** The table summarizes the experimental conditions for cellular kinetic analyses used in this study and previous reports of NOB strains. Cell preparation and analysis conditions (temperature, pH, nitrite concentrations) are indicated.

| Topology of NXR | Strain | Temperature (°C) | Cell preparation | Tested pH | Number of replicates | $K_{m(app)}$ (µM) | $V_{max}$ (µmol N (mg protein)$^{-1}$ h$^{-1}$) | $a^o$ (l (g cells)$^{-1}$ h$^{-1}$) | Reference |
|---|---|---|---|---|---|---|---|---|---|
| cNXR | Strain JJSN | 25 | pH 6.0<br>1 mM $NaNO_2$ | 5.0 | 5 | 11.3 ± 2.7 | 37.2 ± 1.4 | 377.5 ± 86.8 | This study |
| | | | | 7.0 | 3 | 493.7 ± 77.6 | 47.2 ± 2.1 | 12.1 ± 1.1 | |
| | *Nitrobacter winogradksyi* Nb-255 | 28 | pH 7.5<br>1 mM $NaNO_2$ | 5.5 | 5 | 20.4 ± 4.3 | 25.2 ± 1.2 | 178.4 ± 47.9 | This study |
| | | | | 7.5 | 3 | 25.9 ± 5.2 | 68.8 ± 2.5 | 440.5 ± 167.5 | |
| | | | pH 7.5<br>10 mM $NaNO_2$ | 5.5 | 6 | 19.2 ± 1.2 | 76.0 ± 4.0 | 577.1 ± 149.6 | |
| | | | | 7.5 | 3 | 388.0 ± 52.2 | 43.7 ± 1.3 | 14.4 ±2.3 | |
| | | | pH 7.5<br>9 mM $NaNO_2$ | 7.5 | 3 | 309.0 ± 92.0 | 78.0 ± 5.0 | 31.3[b] | ref[1] |
| | | 25 | pH 7.3<br>30 mM $NaNO_2$ | 7.3 | 3 | 36–260 | N/A[a] | N/A[a] | ref[2] |
| | *Nitrobacter vulgaris* DSM 10236 | 28 | pH 7.5<br>9 mM $NaNO_2$ | 7.5 | 3 | 49.0 ± 11.0 | 164.0 ± 9.0 | 415.7[b] | ref[1] |

| | | | | | | | | | |
|---|---|---|---|---|---|---|---|---|---|
| | *Nitrobacter hamburgensis* X14 | 28 | pH 7.5 9 mM $NaNO_2$ | 7.5 | 3 | 544.0 ± 55.0 | 64.0 ± 1.0 | 14.6 [b] | ref[1] |
| | | 25 | pH 7.5 5 mM $(NH_4)_2SO_4$ [C] | 7.5 | 5 | 706–1240 | N/A[a] | N/A[a] | ref[3] |
| | *Nitrobacter sp.* Nb-311A | 28 | pH 7.5 30 mM $NaNO_2$ | 7.5 | 3 | 27.6 ± 6.7 | 95.2 ± 7.0 | 427.4[b] | ref[4] |
| | *'Ca. Nitrobacter laanbroekii'* NHB1 | 25 | pH 6.0 N/A $NaNO_2$ | 6.0 | 4 | 53.5 ± 16.6 | N/A[a] | N/A[a] | ref[5] |
| | *'Ca. Nitrobacter acidiphilus'* ACIDNB-1 | 22 | pH 4.6–5.5 4.2 mM $NaNO_2$ | 6.0 | | 13.6 ± 2.1 | N/A[a] | N/A[a] | ref[6] |
| | *Nitrococcus mobilis* Nb-231 | 28 | pH 7.5 30 mM $NaNO_2$ | 5.0 | 4 | 9.5 ± 0.7 | 92.8 ± 1.5 | 1,042.5 ± 144.6 | This study |
| | | | | 7.0 | 3 | 132.6 ± 19.1 | 59.1 ± 2.7 | 60.6 ± 29.5 | |
| | | | pH 7.5 30 mM $NaNO_2$ | 7.5 | 3 | 119.7 ± 34.0 | 141.0 ± 10.6 | 146.0[b] | ref[4] |
| | *Nitrolancea hollandica* Lb | 37 | pH 6.9–7.4 20 – 50 mM $NaNO_2$ | 6.9 – 7.4 | N/A[a] | 1,000 | N/A[a] | N/A[a] | ref[7] |
| pNXR | *Nitrospira* | 37 | pH 7.5 | 6.0 | 4 | 4.5 ± 1.6 | 17.1 ± 0.8 | 538.9 ± 220.9 | This study |

| | | | | | | | | | |
|---|---|---|---|---|---|---|---|---|---|
| | *moscoviensis* M1 | | 1 mM $NaNO_2$ | 8.0 | 4 | 4.4 ± 1.0 | 21.6 ± 0.7 | 662.7 ± 295.9 | |
| | | | pH 7.5 5.7 mM $NaNO_2$ | 7.5 | 3 | 9.0 ± 3.0 | 18.0 ± 1.0 | 247.8[b] | ref[1] |
| | *'Ca. Nitrospira defluvii'* A17 | 28 | pH 7.5 9 mM $NaNO_2$ | 7.5 | 3 | 9.0 ± 3.0 | 48.0 ± 2.0 | 660.9[b] | ref[1] |
| | *Nitrospira lenta BS10* | 28 | pH 7.5 9 mM $NaNO_2$ | 7.5 | 3 | 27.0 ± 11.0 | 20.0 ± 2.0 | 917.9[b] | ref[1] |
| | *Nitrrospira japonica* NJ1 | 29 | pH 7.8 0.7 mM $NaNO_2$ | 7.8 | 3 | 10.0 ± 2.0 | 31.0 ± 5.0 | 384.1[b] | ref[8] |
| | *Nitrospira* sp. ND1 | 29 | pH 7.8 0.7 mM $NaNO_2$ | 7.8 | 3 | 6.0 ± 1.0 | 45 ± 7 | 929.3[b] | ref[8] |
| | *Nitrospira marina* Ecomares 2.1 | 28 | pH 7.8 0.7 mM $NaNO_2$ | 7.8 | 4 | 54.0 ± 11.9 | 21.4 ± 1.2 | 49.1[b] | ref[4] |
| | '*Ca.* Nitrotoga fabula' KNB | 25 | pH 7.5 1 mM $NaNO_2$ | 6.5 | 2 | 39.4 ± 7.3 | 44.1 ± 2.3 | 144.7 ± 32.3 | This study |
| | | | | 8.0 | 3 | 75.0 ± 10.0 | 48.8 ± 1.4 | 80.9 ± 10.9 | |
| | | | | 7.5 | 4 | 89.3 ± 3.4 | 27.5 ± 7.4 | 54.3 ± 15.5 | ref[9] |
| | '*Ca.* Nitrotoga artica' | 17 | pH 7.8 0.7 mM $NaNO_2$ | 7.5 | 4 | 58.0 ± 28.0 | 20.0 ± 2.0 | 38.3 ± 11.0 | ref[1] |

| | | | | | | | | | |
|---|---|---|---|---|---|---|---|---|---|
| | '*Ca.* Nitrotoga sp.' AM1 | 16 | pH 7.8<br>1.4 mM $NaNO_2$ | 7.8 | 3 | 24.7 ± 9.8 | N/A[a] | N/A[a] | ref[10] |
| | *Nitrospina wastonii* 347 | 28 | pH 7.8<br>5.8 mM $NaNO_2$ | 7.8 | 2 | 18.7 ± 2.1 | 36.8 ± 2.2 | 243.9[b] | ref[4] |

[a] N/A denotes data not available.

[b]The $a^o$ value lacking a standard deviation (s.d.) was calculated from average $K_{m(app)}$ and $V_{max}$ values, due to the absence of individual replicate data.

[c]Cells were prepared from coculture with *Nitrosomonas europaea,* an ammonia-oxidizing bacterium.

**Supplementary Table 4. List of candidate nitrite/nitrate transport proteins identified in selected NOB genomes.** Listed are the GenBank accession IDs for each analyzed genome, NCBI-designated organism names, and the corresponding NXR topology type (cNXR or pNXR) for each strain. Protein accession IDs for candidate transporters are provided. When multiple homologs are identified within a genome, entries are separated by line breaks within the same data field.

| **Topology of NXR** | **Strain** | **Formate/Nitrite transporter family** | | **Nitrite/Nitrate porter** | | | **ABC transport** |
|---|---|---|---|---|---|---|---|
| | | **NirC (PF01226)** | **FocA (PF01226)** | **NarK-n (PF7690)** | **NarK-c (PF7690)** | **NrtP (PF07690)** | **NrtA (cd13553)** |
| cNXR | Strain JJSN | JJSN_00253 | | **JJSN_00145**[a]<br>JJSN_02717<br>JJSN_00617<br>JJSN_00593 | JJSN_01191 | | |
| | '*Ca.* Nitrobacter acidophilus' ACIDNB-1 (GCF_033852255.1) | WP_319797425.1,<br>WP_319798015.1 | | **WP_292617488.1**[a],<br>WP_292695622.1,<br>WP_319796629.1 | WP_319798838.1 | | |
| | *Nitrobacter winogradskyi* Nb-255 (GCA_000012725.1) | ABA06256.1 | | **ABA04044.1**[a] | ABA04680.1 | | ABA03724.1 |
| | *Nitrobacter hamburgensis* X14 (GCA_000013885.1) | ABE61919.1 | | **ABE64173.1**[a],<br>ABE62266.1 | ABE63249.1 | | |
| | *Nitrobacter vulgaris* DSM 10236 (GCA_031453995.1)" | MDR6304105.1 | | **MDR6306289.1**[a],<br>MDR6302966.1,<br>MDR6306204.1 | MDR6303946.1 | | MDR6306110.1 |
| | '*Ca*. Nitrobacter laanbroekii' NHB1 (GCA_036964665.1) | MEH6951354.1 | | **MEH6952355.1**[a],<br>MEH6951011.1,<br>MEH6952691.1 | MEH6950278.1,<br>MEH6952825.1 | | |
| | *Nitrobacter* sp. Nb-311A (GCA_000152905.1) | EAQ35807.1 | | **EAQ33961.1**[a],<br>EAQ36645.1,<br>EAQ36371.1 | EAQ34794.1 | | |

| | | | | | | |
|---|---|---|---|---|---|---|
| | *Nitrobacter* sp. TKz-YC01 (GCA_047824785.1) | XOP68878.1 | **XOP69876.1**[a], XOP68471.1, XOP70626.1 | XOP70415.1 | | XOP69535.1 |
| | *Nitrobacter* sp. AFB05 (GCA_047149785.1) | GAB1716918.1, GAB1718218.1 | **GAB1714970.1**[a], GAB1716555.1 | GAB1716010.1 | | |
| | *Nitrobacter* sp. ECS31B4 (GCA_025698885.1) | MCV0386368.1 | **MCV0387771.1**[a], MCV0386021.1 | MCV0387649.1 | | |
| | *Nitrobacter* sp. HKST-UBA79 (GCA_020441525.1) | MCB1393297.1 | **MCB1393177.1**[a] | MCB1394038.1 | | |
| | *Nitrococcus mobilis* Nb-231 (GCA_000153205.1) | EAR22762.1 | EAR22036.1 | EAR20979.1 | | |
| | *Nitrolancea hollandica* Lb (GCA_000297255.1) | CCF84986.1 | CCF85210.1 | | | |
| pNXR | '*Ca*. Nitrospira allomarina' VA (GCA_032050975.1) | WNM58070.1 | WNM57171.1 | | | |
| | *Nitrospira defluvii* ZN2 (GCA_905220995.1) | CAE6737653.1, CAE6745279.1 | CAE6736094.1 | | | |
| | *Nitrospira japonica* NSJP_Ch1 (GCA_900169565.1) | SLM50047.1 | SLM48610.1 | | | SLM48575.1 |
| | *Nitrospira lenta* BS10 (GCA_900403705.1) | SPP64249.1, SPP66061.1 | SPP66726.1 | | | SPP64196.1 |
| | *Nitrospira moscoviensis* M-1 (GCA_001273775.1) | | ALA57724.1 | | | ALA57699.1 |
| | '*Ca*. Nitrospira neomarina' DK (GCA_032051675.1) | WNM60213.1 | WNM63201.1 | | | |
| | *Nitrospira tepida* DNF (GCA_947241125.1) | CAI4030805.1 | CAI4031305.1 | | | CAI4031284.1 |
| | *Nitrospina watsonii* 347 (GCA_946900835.1) | CAI2717024.1 | | | CAI2719027.1 | |

| | | |
|---|---|---|
| *Nitrospina gracilis* 3/211 (GCA_000341545.2) | CCQ89743.1 | CCQ90797.1 |
| '*Ca*. Nitrotoga artica' 6680 (GCA_918378365.1) | CAG9932624.1 | |
| '*Ca* Nitrotoga fabula' ZN8 (GCA_905221025.1) | CAE6690416.1 | |

[a]Associated with the NXR cluster

**Supplementary Table 5. NOB strains and experimental conditions used for nitrite oxidation in this study.**

| Topology of NXR | Strain | Growth condition | | | Medium | Isolation source | Reference |
|---|---|---|---|---|---|---|---|
| | | pH | Temperature (°C) | Nitrite (µM) | | | |
| cNXR | Strain JJSN | 4.0–9.0 | 25 | 10–1,000 | DSMZ 756c | Forest soil | This study |
| | *Nitrobacter winogradskyi* Nb-255 | 5.0–9.0 | 28 | 10–10,000 | DSMZ 756c | Soil | This study and Nowka *et al*[1] |
| | *Nitrococcus mobilis* Nb-231 | 5.0–9.0 | 28 | 10–30,000 | Seawater medium | Marine | This study and Watson *et al*[11] |
| pNXR | *Nitrospira moscoviensis* M1 | 5.0–9.0 | 37 | 10–1,000 | DSMZ 756c | Heating system | This study and Nowka *et al*[1] |
| | ‘*Ca.* Nitrotoga fabula’ KNB | 5.0–9.0 | 25 | 10–1,000 | DSMZ 756c | Activated sludge | This study and Kitzinger *et al*[9] |

**Supplementary Table 6. Primer sets used for the RT-qPCR analysis of gene expression in *N. winogradskyi* Nb-255.**

| Primer (Forward and Reverse) | Description (Accession number) | Sequence (5′ to 3′) | Length (bp) | Tm (℃) |
|---|---|---|---|---|
| Nwi_recA_763F<br>Nwi_recA_981R | Recombinase subunit alpha (*recA*)<br>(ABA04099.1; Nwi_0834) | GTC AAG GTC GTG AAG AA<br>GGT GAT GTC AGG ATT GG | 218 | 55 |
| Nwi_nxrA_1887F<br>Nwi_nxrA_2133R | Nitrite oxidoreductase subunit alpha (*nxrA*)<br>(ABA04039.1; Nwi_0774) | CAC TCC GCA CTT TAA TC<br>GTG AGC ACC AAG ATA GT | 246 | 55 |
| Nwi_nriC_397F<br>Nwi_nirC_617R | formate/nitrite transporter (*nirC*)<br>(ABA06256.1; Nwi_3006) | TTC TCA CTA ACC AAG GG<br>ATC GGT ATC CAC ATC AG | 220 | 55 |
| Nwi_narKn_1008F<br>Nwi_narKn_1209R | Nitrate/Nitrite porter (*narK*-n) in NXR operon<br>(ABA04044.1; Nwi_0779) | GCT GAT CCC GAT CTA TT<br>ATC TCT AAA GCC CAC TC | 201 | 55 |
| Nwi_narKc_682F<br>Nwi_narKc_869R | Nitrate/Nitrite porter (*narK*-c)<br>(ABA04680.1; Nwi_1419) | CCG CAA TAC TTC AAG AC<br>GGA TAG GAC AGC AGA AA | 187 | 55 |

### Supplementary Figure 1–12

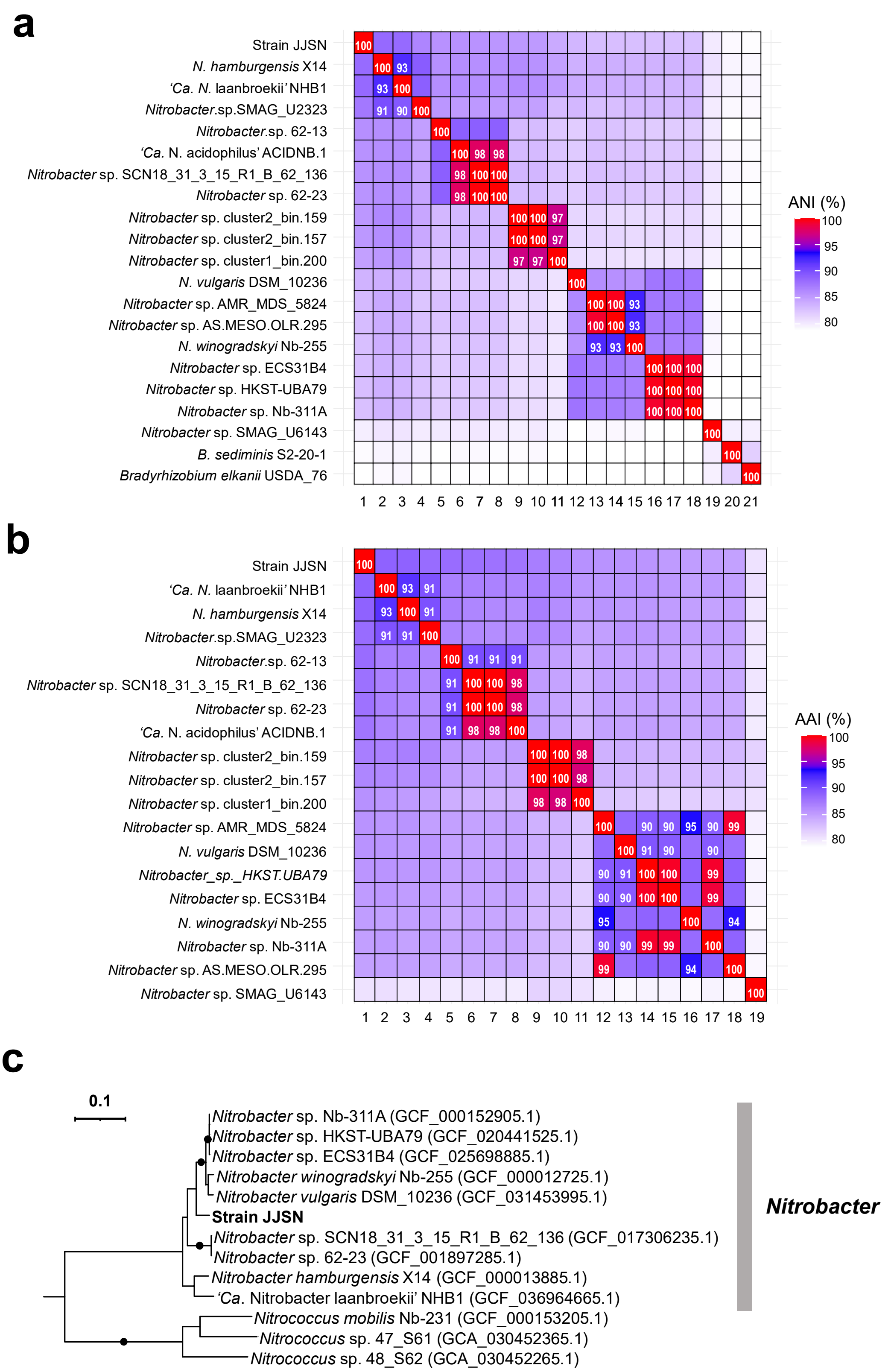

**Supplementary Figure 1. Comparative ANI and AAI values and NxrA phylogenetic tree of *Nitrobacter* strains. a,** Pairwise average nucleotide identity (ANI) values among *Nitrobacteraceae* family members. **b,** Pairwise average amino acid identity (AAI) values for the same *Nitrobacter* strains. White numbers within boxes in panels **a** and **b** indicate pairwise identity values exceeding 90%. **c,** Maximum-likelihood tree of *Nitrobacter* NxrA amino acid sequences that are associated with NXR gene clusters. *Nitrococcus* NxrA amino acid sequences were used as an outgroup. Bracketed numbers indicate NCBI genome accession numbers. The strain obtained in this study is shown in bold. Branch support values ≥ 95% are indicated by black circles.

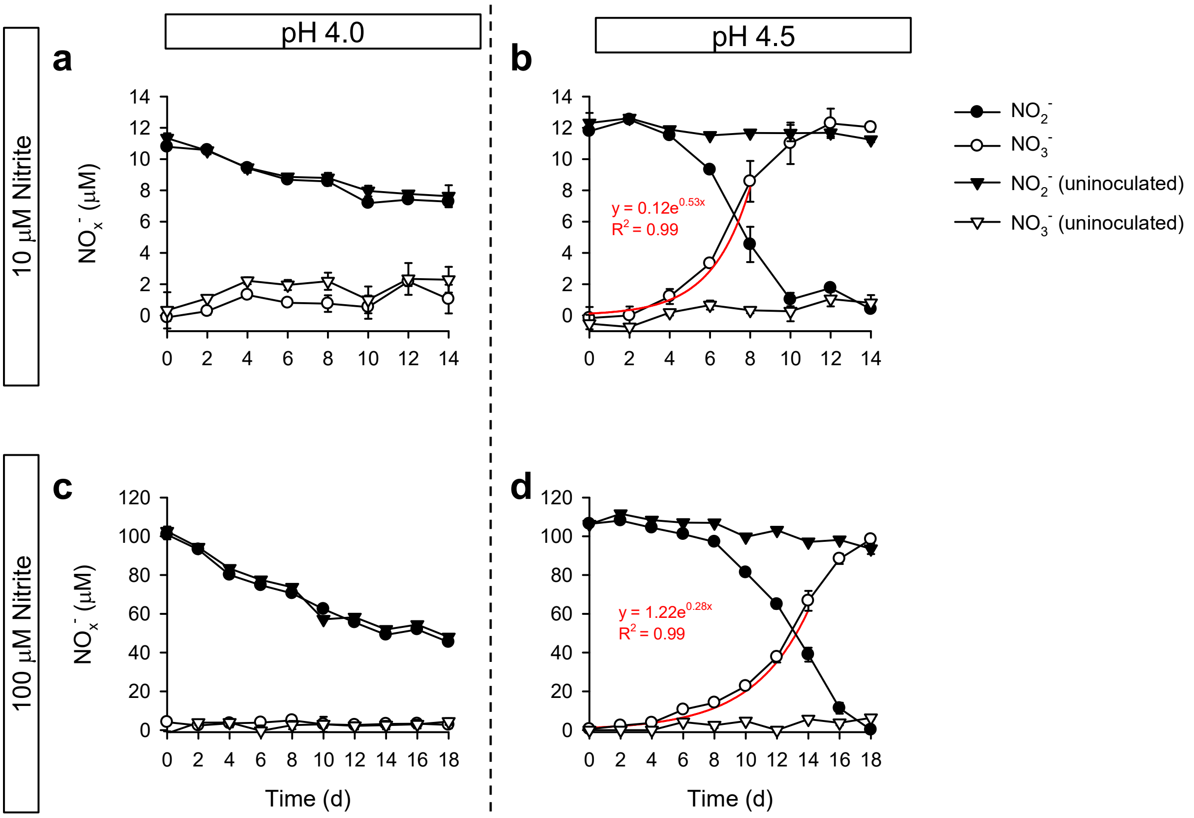


**Supplementary Figure 2. Nitrite oxidation activity of strain JJSN under acidic conditions.** Nitrite and nitrate concentrations in cultures of strain JJSN at pH 4.0 (**a, c**) and 4.5 (**b, d**), with two initial nitrite concentrations (10 and 100 µM). Abiotic controls (uninoculated medium) were included for comparison. Red curves represent exponential fits of nitrate accumulation in biological cultures, corrected for abiotic nitrite decomposition. Data are presented as mean ± s.d. ($n$ = 3).

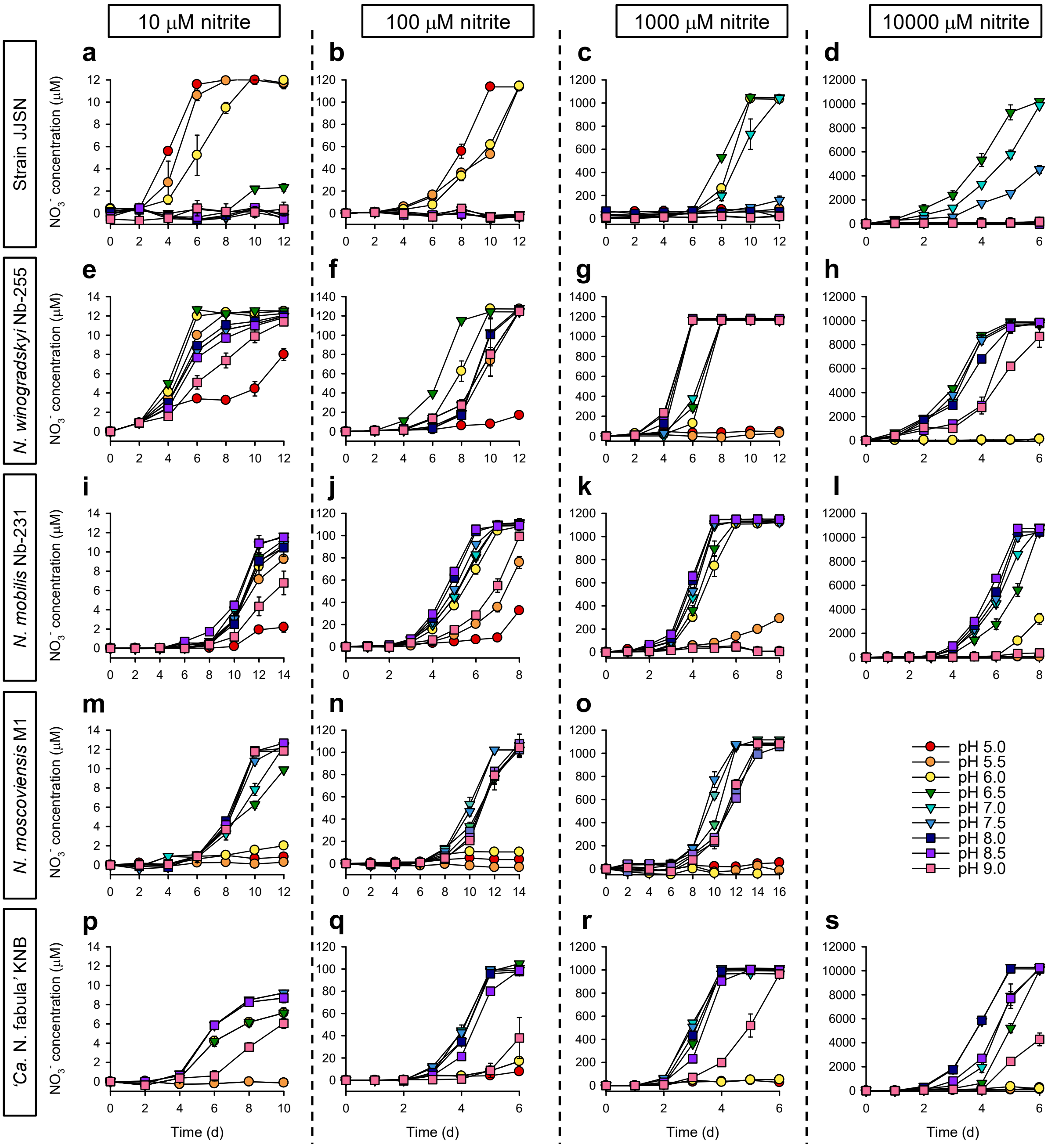


**Supplementary Figure 3. Nitrite oxidation profiles under varying pH and nitrite concentrations.** Nitrate production under four nitrite concentrations (10–10,000 μM) using strain JJSN (**a**–**d**), *N. winogradskyi* Nb-255 (**e**–**h**), *N. mobilis* Nb-231 (**i**–**l**), *N. moscoviensis* M1 (**m**–**o**), and "*Ca*. N. fabula" KNB (**p**–**s**). Data are presented as mean ± s.d. ($n$ = 3).

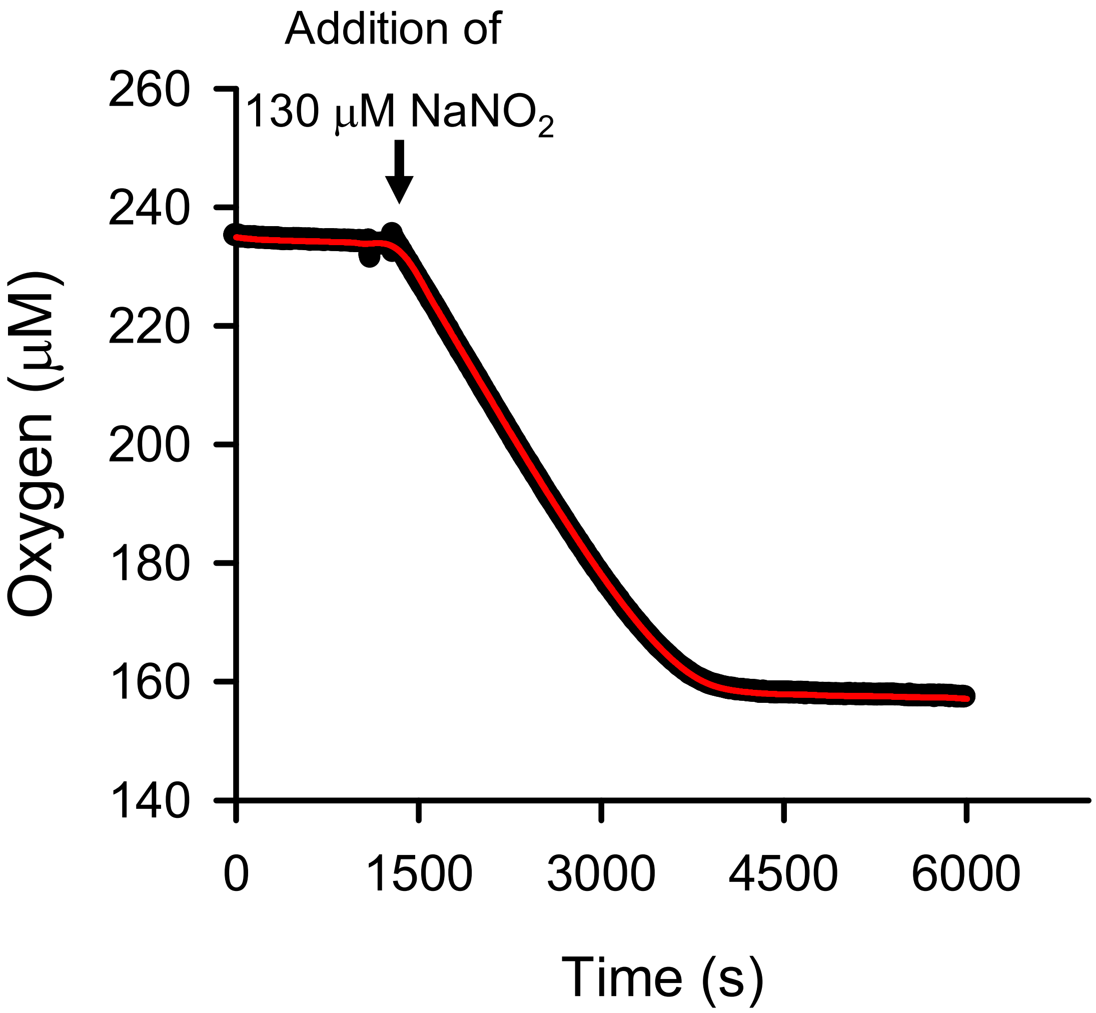


**Supplementary Figure 4. Nitrite-dependent stoichiometric oxygen uptake by cells of strain JJSN.** Mid-log-phase cells were harvested and incubated at 25 °C and pH 5.0. A stable background oxygen uptake rate for approximately 0.5 h was observed before nitrite addition (130 µM $NaNO_2$). The red line represents smoothed data processed using the 'Smooth 2D data' function in SigmaPlot 10.0 (SPSS Inc., USA).

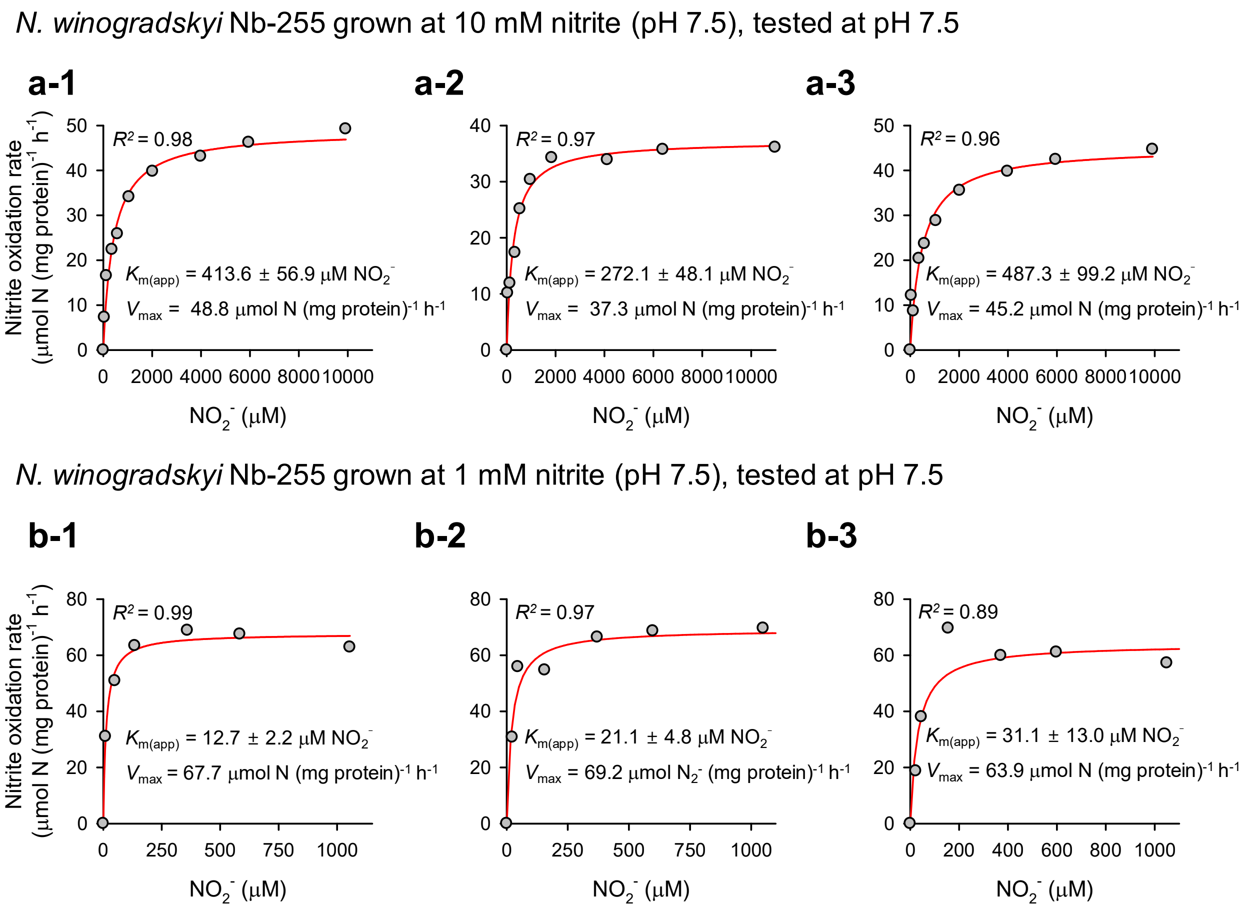

N. winogradskyi Nb-255 grown at 10 mM nitrite (pH 7.5), tested at pH 7.5
a-1
a-2
a-3
Nitrite oxidation rate
(µmol N (mg protein)$^{-1}$ h$^{-1}$)
$R^2$ = 0.98
$K_{m(app)}$ = 413.6 ± 56.9 µM $NO_2^-$
$V_{max}$ = 48.8 µmol N (mg protein)$^{-1}$ h$^{-1}$
$R^2$ = 0.97
$K_{m(app)}$ = 272.1 ± 48.1 µM $NO_2^-$
$V_{max}$ = 37.3 µmol N (mg protein)$^{-1}$ h$^{-1}$
$R^2$ = 0.96
$K_{m(app)}$ = 487.3 ± 99.2 µM $NO_2^-$
$V_{max}$ = 45.2 µmol N (mg protein)$^{-1}$ h$^{-1}$
$NO_2^-$ (µM)
N. winogradskyi Nb-255 grown at 1 mM nitrite (pH 7.5), tested at pH 7.5
b-1
b-2
b-3
$R^2$ = 0.99
$K_{m(app)}$ = 12.7 ± 2.2 µM $NO_2^-$
$V_{max}$ = 67.7 µmol N (mg protein)$^{-1}$ h$^{-1}$
$R^2$ = 0.97
$K_{m(app)}$ = 21.1 ± 4.8 µM $NO_2^-$
$V_{max}$ = 69.2 µmol $N_2^-$ (mg protein)$^{-1}$ h$^{-1}$
$R^2$ = 0.89
$K_{m(app)}$ = 31.1 ± 13.0 µM $NO_2^-$
$V_{max}$ = 63.9 µmol N (mg protein)$^{-1}$ h$^{-1}$

*N. winogradskyi* Nb-255 grown at 10 mM nitrite (pH 7.5), tested at pH 5.5

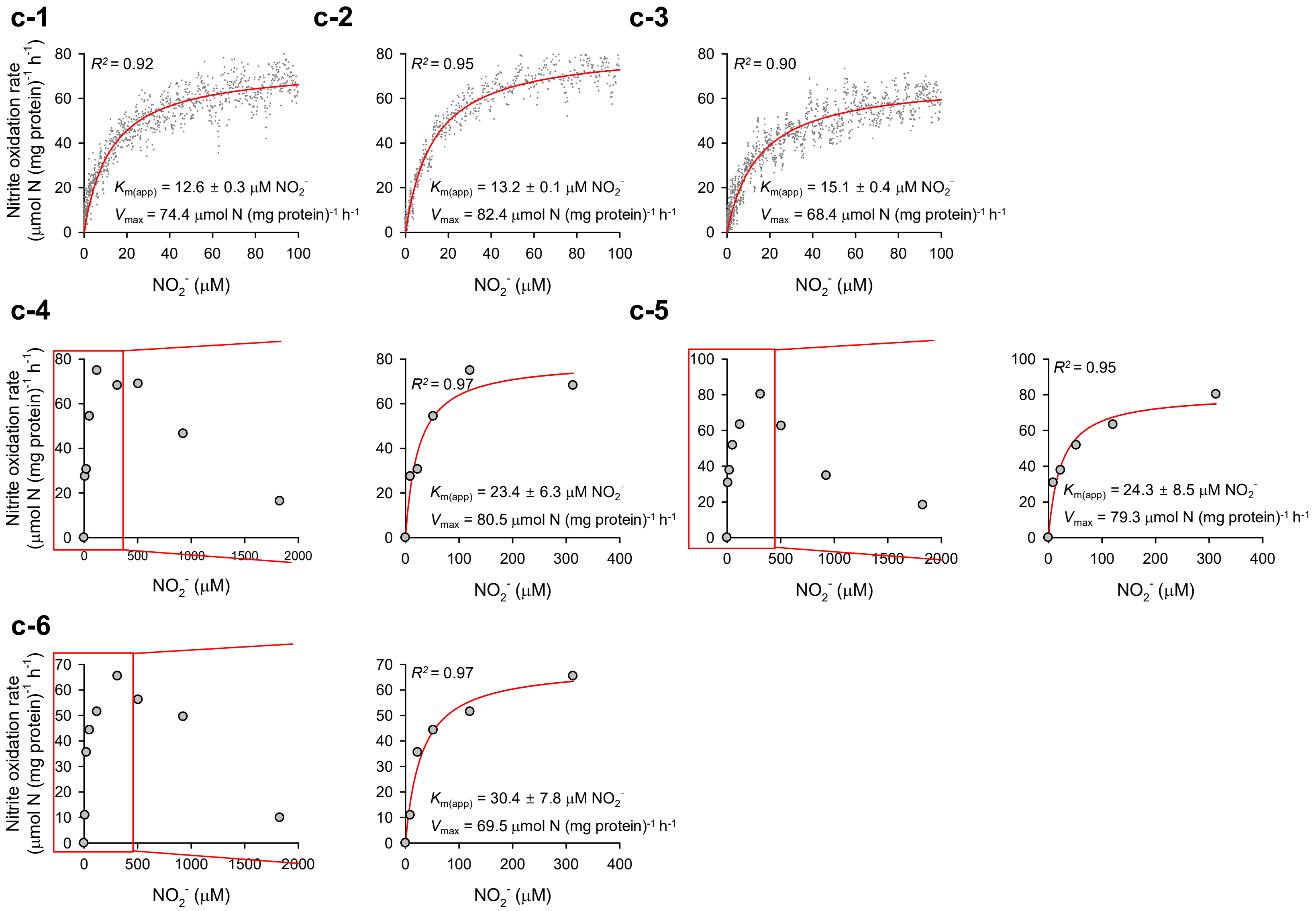

*N. winogradskyi* Nb-255 grown at 1 mM nitrite (pH 7.5), tested at pH 5.5

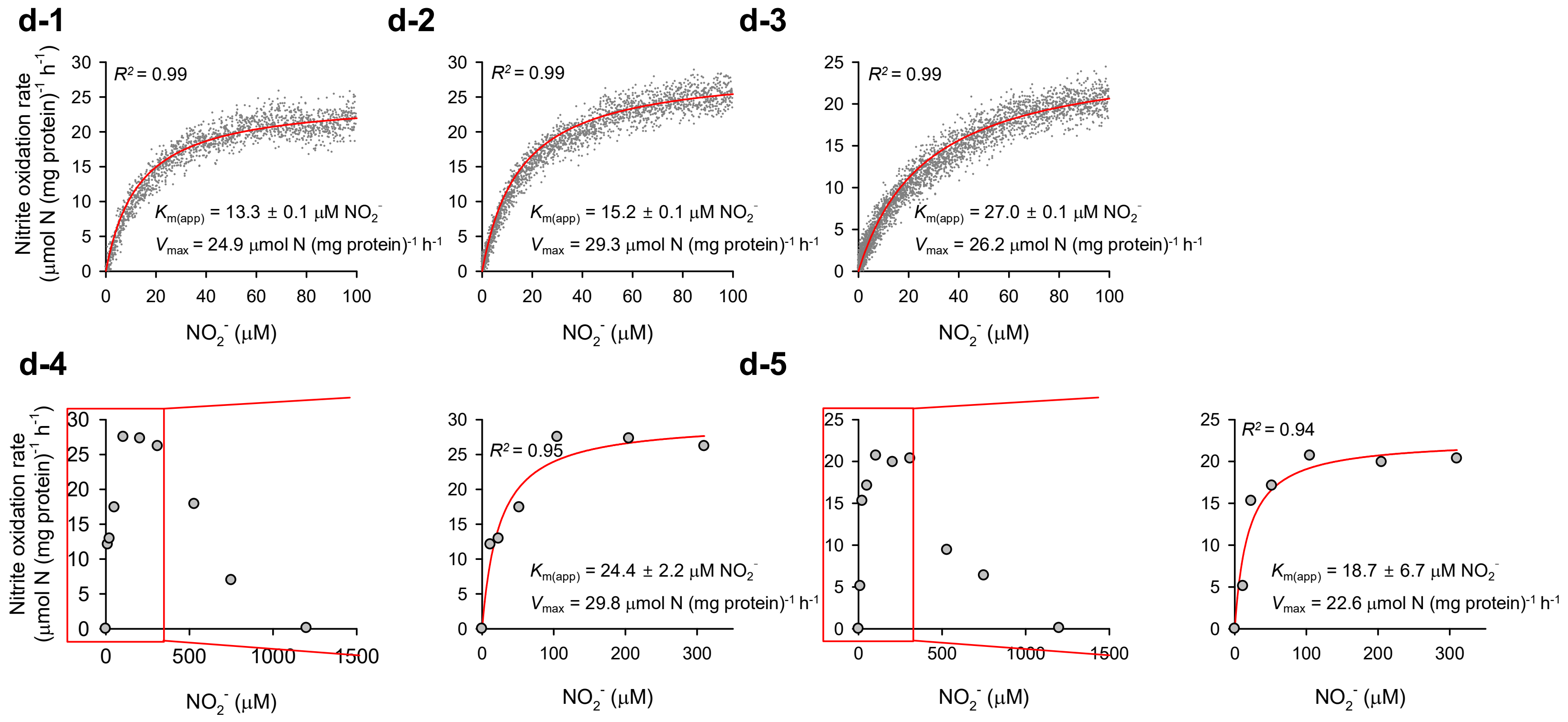

Strain JJSN grown at 1 mM nitrite (pH 6.0), tested at pH 7.0

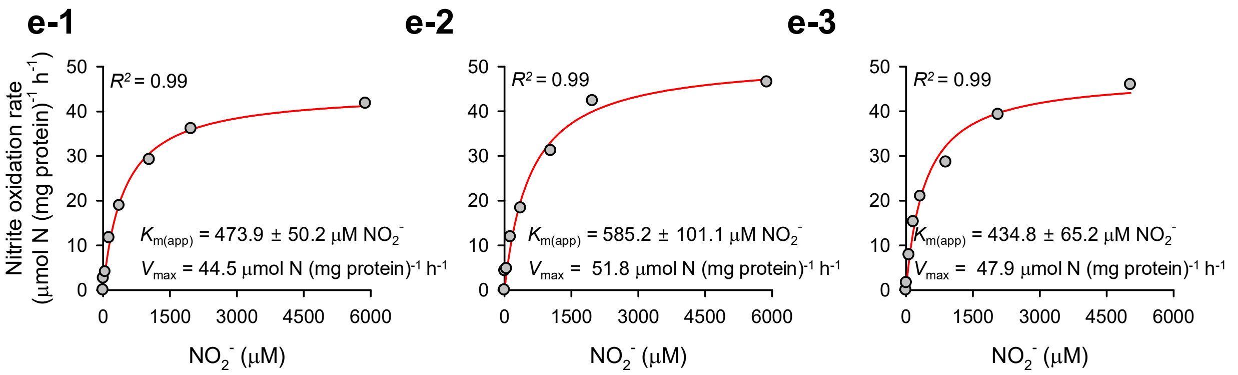


Strain JJSN grown at 1 mM nitrite (pH 6.0), tested at pH 5.0

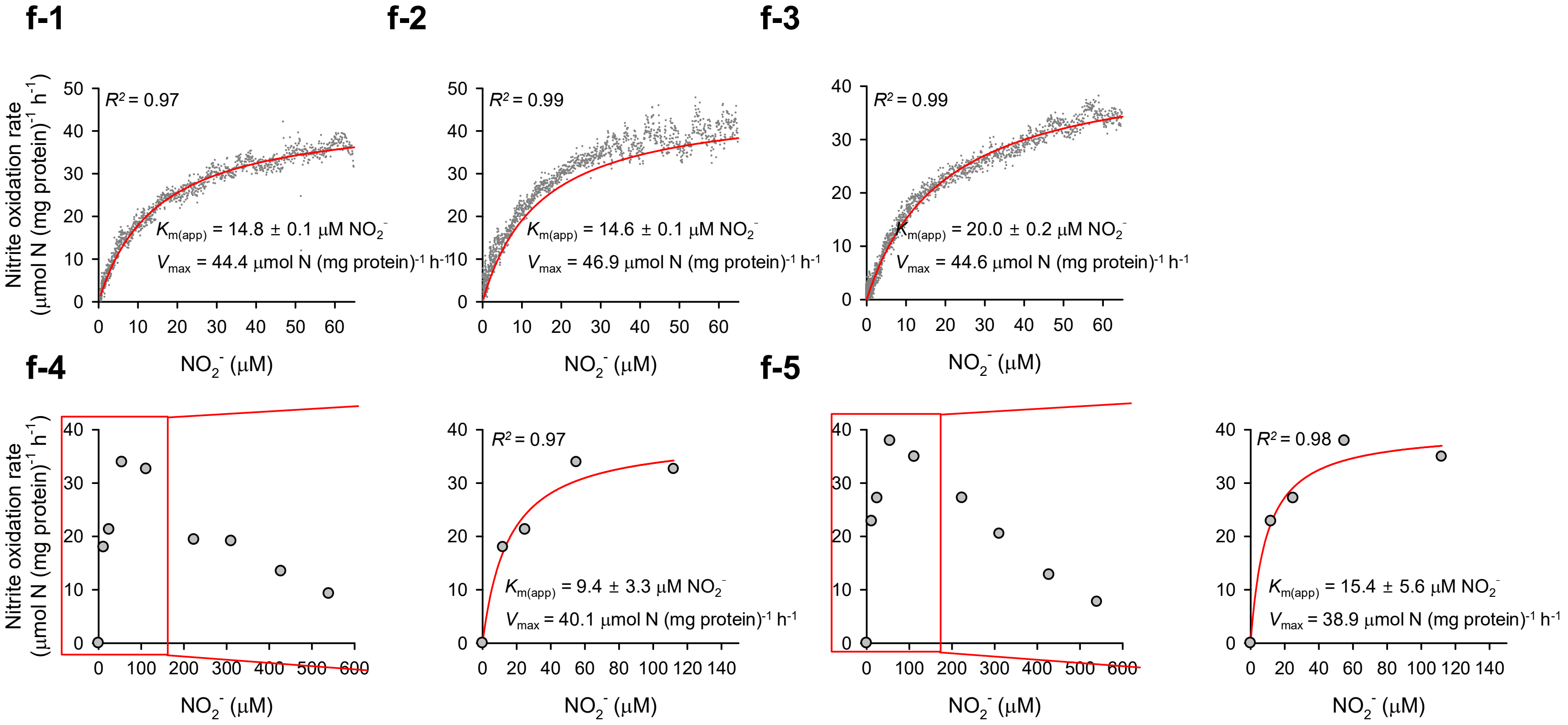

*N. mobilis* Nb-231 grown at 30 mM nitrite (pH 7.5), tested at pH 7.5

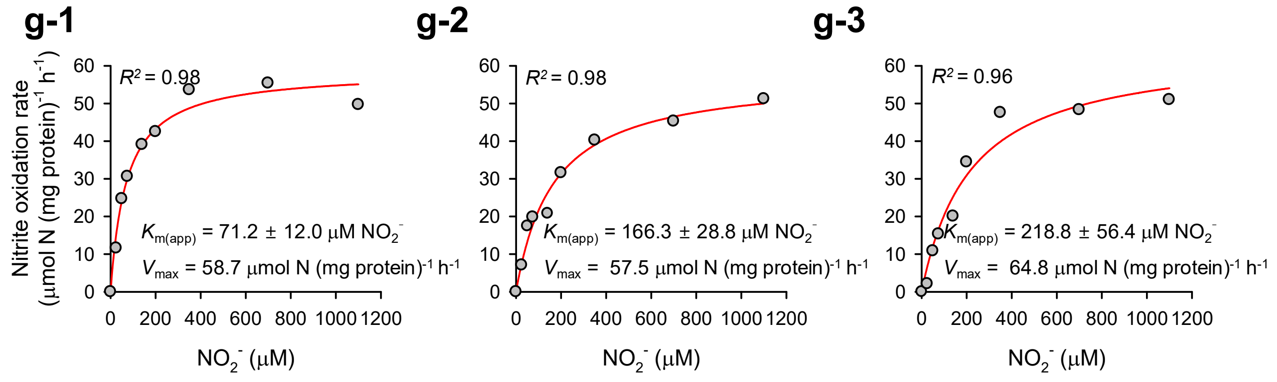


*N. mobilis* Nb-231 grown at 30 mM nitrite (pH 7.5), tested at pH 5.5

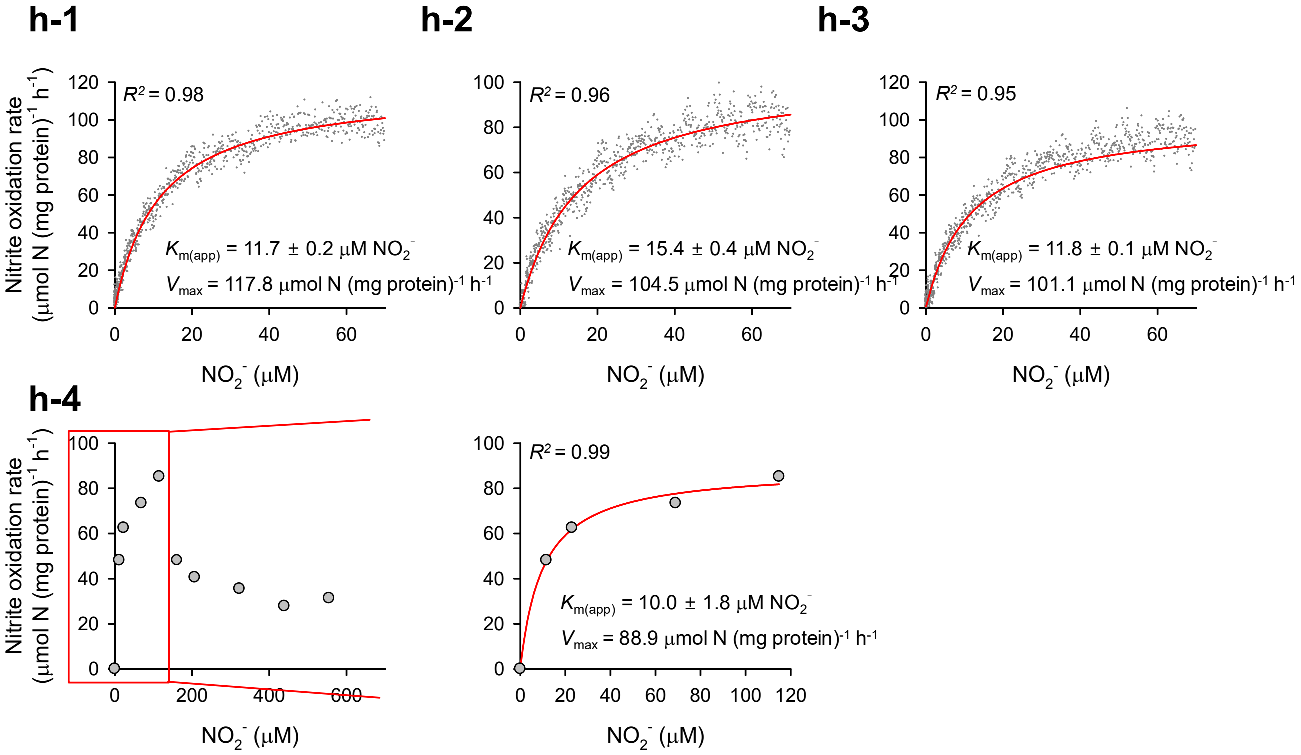

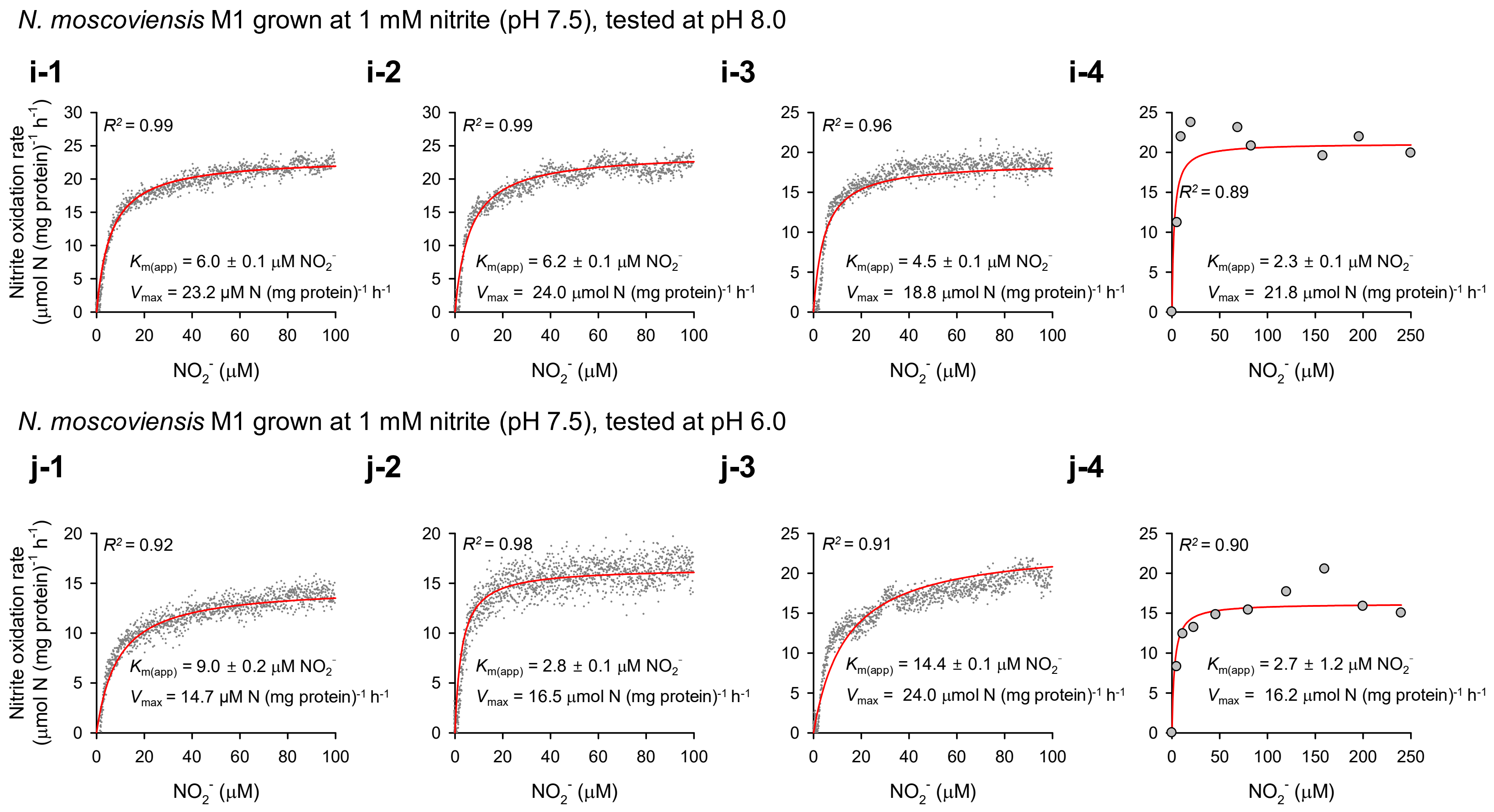
N. moscoviensis M1 grown at 1 mM nitrite (pH 7.5), tested at pH 8.0
i-1
R2 = 0.99
Km(app) = 6.0 ± 0.1 µM NO2-
Vmax = 23.2 µM N (mg protein)-1 h-1
i-2
R2 = 0.99
Km(app) = 6.2 ± 0.1 µM NO2-
Vmax = 24.0 µmol N (mg protein)-1 h-1
i-3
R2 = 0.96
Km(app) = 4.5 ± 0.1 µM NO2-
Vmax = 18.8 µmol N (mg protein)-1 h-1
i-4
R2 = 0.89
Km(app) = 2.3 ± 0.1 µM NO2-
Vmax = 21.8 µmol N (mg protein)-1 h-1
N. moscoviensis M1 grown at 1 mM nitrite (pH 7.5), tested at pH 6.0
j-1
R2 = 0.92
Km(app) = 9.0 ± 0.2 µM NO2-
Vmax = 14.7 µM N (mg protein)-1 h-1
j-2
R2 = 0.98
Km(app) = 2.8 ± 0.1 µM NO2-
Vmax = 16.5 µmol N (mg protein)-1 h-1
j-3
R2 = 0.91
Km(app) = 14.4 ± 0.1 µM NO2-
Vmax = 24.0 µmol N (mg protein)-1 h-1
j-4
R2 = 0.90
Km(app) = 2.7 ± 1.2 µM NO2-
Vmax = 16.2 µmol N (mg protein)-1 h-1
Nitrite oxidation rate (µmol N (mg protein)-1 h-1)
NO2- (µM)

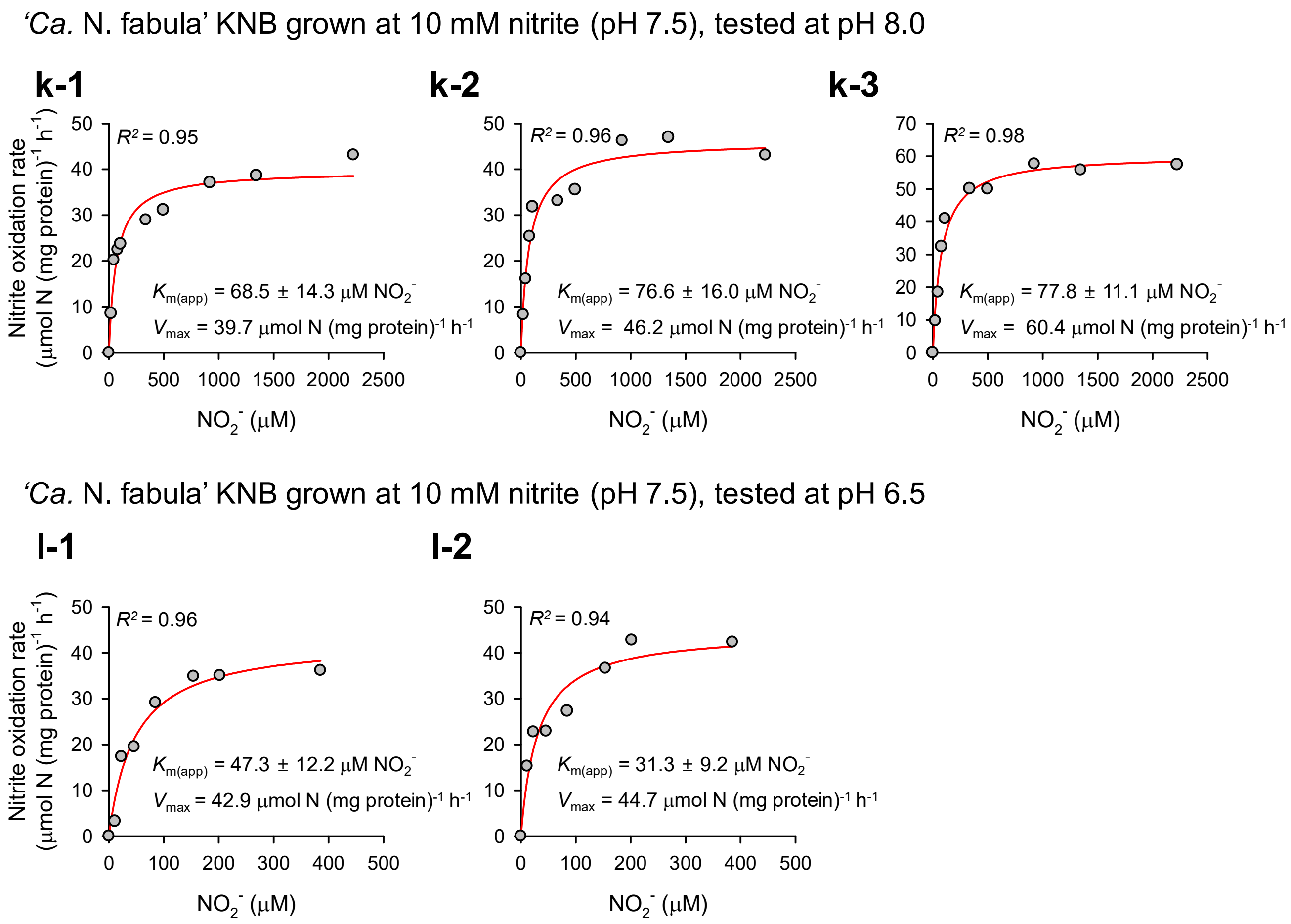
'Ca. N. fabula' KNB grown at 10 mM nitrite (pH 7.5), tested at pH 8.0
k-1
R² = 0.95
Km(app) = 68.5 ± 14.3 µM NO2-
Vmax = 39.7 µmol N (mg protein)-1 h-1
k-2
R² = 0.96
Km(app) = 76.6 ± 16.0 µM NO2-
Vmax = 46.2 µmol N (mg protein)-1 h-1
k-3
R² = 0.98
Km(app) = 77.8 ± 11.1 µM NO2-
Vmax = 60.4 µmol N (mg protein)-1 h-1
Nitrite oxidation rate (µmol N (mg protein)-1 h-1)
NO2- (µM)
'Ca. N. fabula' KNB grown at 10 mM nitrite (pH 7.5), tested at pH 6.5
l-1
R² = 0.96
Km(app) = 47.3 ± 12.2 µM NO2-
Vmax = 42.9 µmol N (mg protein)-1 h-1
l-2
R² = 0.94
Km(app) = 31.3 ± 9.2 µM NO2-
Vmax = 44.7 µmol N (mg protein)-1 h-1

**Supplementary Figure 5. Nitrite oxidation kinetics of five NOB strains.** Michaelis–Menten plots for *N. winogradskyi* Nb-255 (**a**–**d**), strain JJSN (**e**, **f**), *N. mobilis* Nb-231 (**g**, **h**), *N. moscoviensis* M1 (**i**, **j**), and '*Ca*. N. fabula' (**k**, **l**). Nitrite oxidation rates from microsensor measurements of substrate-dependent $O_2$ consumption, using either multiple discrete slopes and/or a single continuous trace. Only discrete slopes at non-inhibitory nitrite concentrations (highlighted with red boxes in each panel) were used to calculate kinetic parameters. Apparent half-saturation constants ($K_{m(app)}$) and maximum oxidation rates ($V_{max}$) were obtained from fitting the data to the Michaelis–Menten equation. The red line indicates the best-fit curve. Standard deviations of the kinetic estimates derived from non-linear regression are reported. Microrespirometry conditions and the number of biological replicates are detailed in **Supplementary Table 3**.

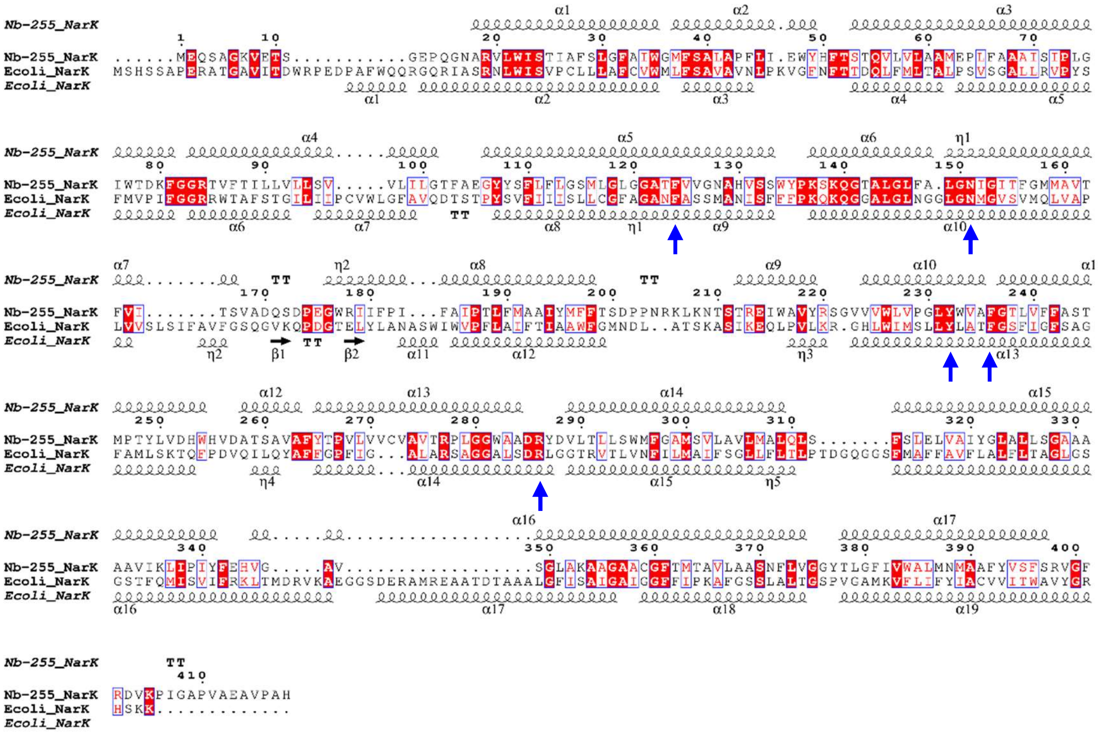


**Supplementary Figure 6. Multiple sequence alignment of NarK proteins.** NarK amino acid sequences from *N. winogradskyi* Nb-255 (Nb-255_NarK-c; ABA04044.1**)** and *Escherichia coli* (Ecoli_NarK; PDB-ID: 4IU9) were aligned. The alignment highlights conserved residues and structural features. Identical residues are shown in white on a red background, and similar residues in red on a white background. Predicted secondary structure of Nb-255_NarK based on homology modeling is shown above the alignment; structural elements derived from the crystal structure of Ecoli_NarK are shown below. α-helices (α1–α17) are indicated by coiled symbols, β-strands (β) by black arrows, and $3_{10}$-helices (η) and β-turns (TT) by text labels. Conserved residues involved in nitrite/nitrate binding (F124, N151, Y232, F236, and R277) are marked with blue vertical arrows.

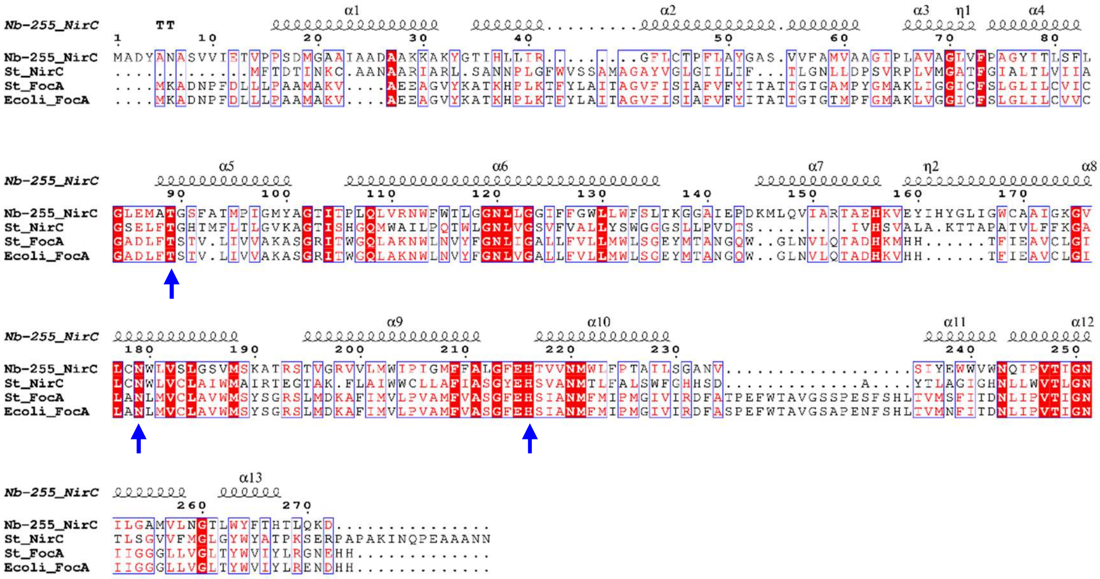


**Supplementary Figure 7. Multiple sequence alignment of FNT family proteins including NirC and FocA.** NirC amino acid sequences from *N. winogradskyi* Nb-255 (Nb-255_NirC; ABA06256.1) and *Salmonella typhimurium* (St_NirC; PDB-ID: 4F4C) and FocA sequences from *S. typhimurium* (St_FocA; PDB-ID: 3Q7K) and *Escherichia coli* (Ecoli_FocA; PDB-ID: 3KCU) were aligned. Sequence conservation and functionally essential residues were highlighted. Identical residues are shown in white on a red background, similar residues in red on a white background. The predicted secondary structure of Nb-255_NirC is shown above the alignment. α-helices (α1–α13) are indicated by coiled symbols and β-turns (TT) by text labels. Conserved residues T89, N179, and H216, which are key residues of the transport pathway, are marked by blue vertical arrows.

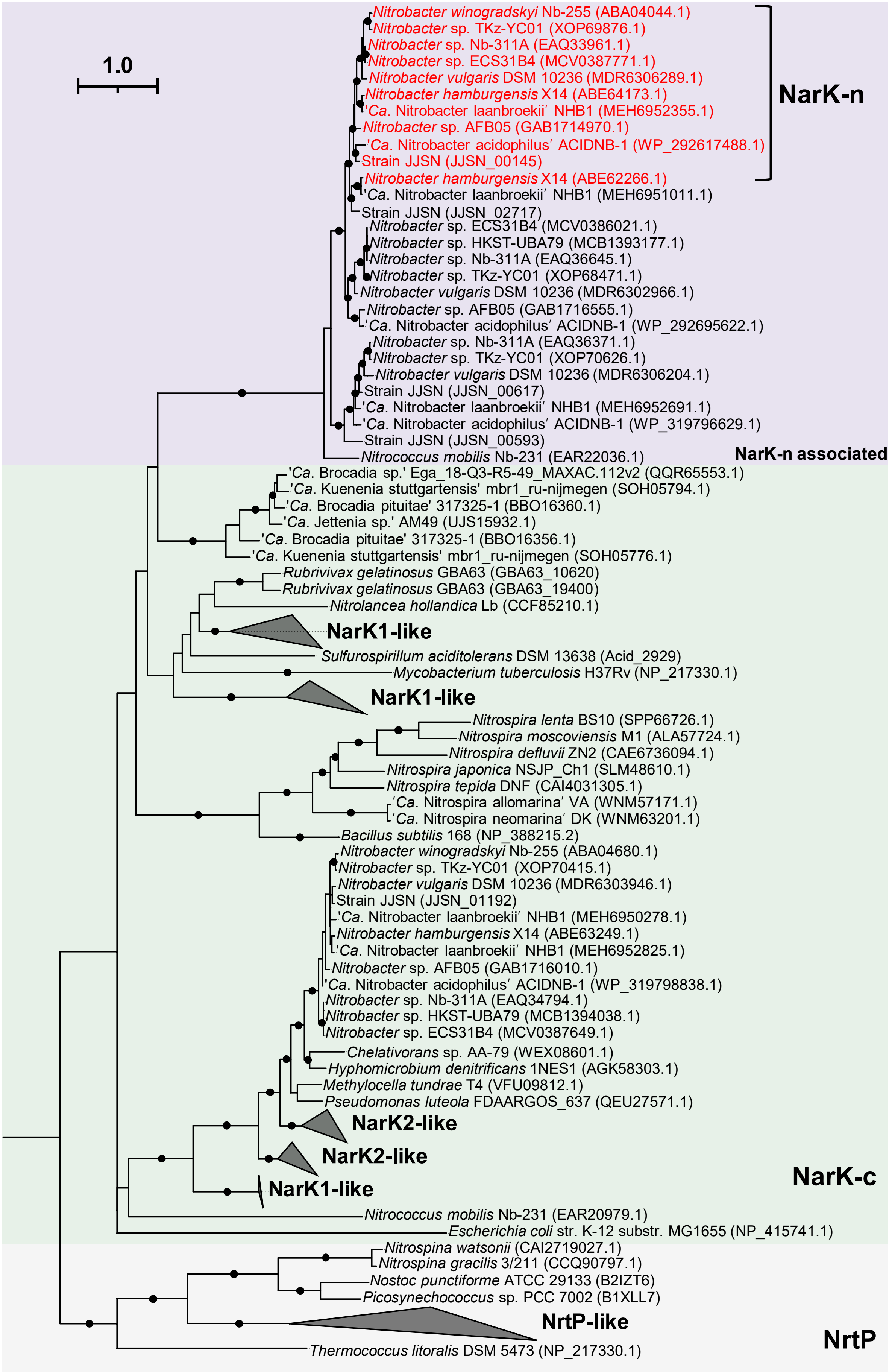
1.0
NarK-n
Nitrobacter winogradskyi Nb-255 (ABA04044.1)
Nitrobacter sp. TKz-YC01 (XOP69876.1)
Nitrobacter sp. Nb-311A (EAQ33961.1)
Nitrobacter sp. ECS31B4 (MCV0387771.1)
Nitrobacter vulgaris DSM 10236 (MDR6306289.1)
Nitrobacter hamburgensis X14 (ABE64173.1)
'Ca. Nitrobacter laanbroekii' NHB1 (MEH6952355.1)
Nitrobacter sp. AFB05 (GAB1714970.1)
'Ca. Nitrobacter acidophilus' ACIDNB-1 (WP_292617488.1)
Strain JJSN (JJSN_00145)
Nitrobacter hamburgensis X14 (ABE62266.1)
'Ca. Nitrobacter laanbroekii' NHB1 (MEH6951011.1)
Strain JJSN (JJSN_02717)
Nitrobacter sp. ECS31B4 (MCV0386021.1)
Nitrobacter sp. HKST-UBA79 (MCB1393177.1)
Nitrobacter sp. Nb-311A (EAQ36645.1)
Nitrobacter sp. TKz-YC01 (XOP68471.1)
Nitrobacter vulgaris DSM 10236 (MDR6302966.1)
Nitrobacter sp. AFB05 (GAB1716555.1)
'Ca. Nitrobacter acidophilus' ACIDNB-1 (WP_292695622.1)
Nitrobacter sp. Nb-311A (EAQ36371.1)
Nitrobacter sp. TKz-YC01 (XOP70626.1)
Nitrobacter vulgaris DSM 10236 (MDR6306204.1)
Strain JJSN (JJSN_00617)
'Ca. Nitrobacter laanbroekii' NHB1 (MEH6952691.1)
'Ca. Nitrobacter acidophilus' ACIDNB-1 (WP_319796629.1)
Strain JJSN (JJSN_00593)
Nitrococcus mobilis Nb-231 (EAR22036.1)
NarK-n associated
'Ca. Brocadia sp.' Ega_18-Q3-R5-49_MAXAC.112v2 (QQR65553.1)
'Ca. Kuenenia stuttgartensis' mbr1_ru-nijmegen (SOH05794.1)
'Ca. Brocadia pituitae' 317325-1 (BBO16360.1)
'Ca. Jettenia sp.' AM49 (UJS15932.1)
'Ca. Brocadia pituitae' 317325-1 (BBO16356.1)
'Ca. Kuenenia stuttgartensis' mbr1_ru-nijmegen (SOH05776.1)
Rubrivivax gelatinosus GBA63 (GBA63_10620)
Rubrivivax gelatinosus GBA63 (GBA63_19400)
Nitrolancea hollandica Lb (CCF85210.1)
NarK1-like
Sulfurospirillum aciditolerans DSM 13638 (Acid_2929)
Mycobacterium tuberculosis H37Rv (NP_217330.1)
NarK1-like
Nitrospira lenta BS10 (SPP66726.1)
Nitrospira moscoviensis M1 (ALA57724.1)
Nitrospira defluvii ZN2 (CAE6736094.1)
Nitrospira japonica NSJP_Ch1 (SLM48610.1)
Nitrospira tepida DNF (CAI4031305.1)
'Ca. Nitrospira allomarina' VA (WNM57171.1)
'Ca. Nitrospira neomarina' DK (WNM63201.1)
Bacillus subtilis 168 (NP_388215.2)
Nitrobacter winogradskyi Nb-255 (ABA04680.1)
Nitrobacter sp. TKz-YC01 (XOP70415.1)
Nitrobacter vulgaris DSM 10236 (MDR6303946.1)
Strain JJSN (JJSN_01192)
'Ca. Nitrobacter laanbroekii' NHB1 (MEH6950278.1)
Nitrobacter hamburgensis X14 (ABE63249.1)
'Ca. Nitrobacter laanbroekii' NHB1 (MEH6952825.1)
Nitrobacter sp. AFB05 (GAB1716010.1)
'Ca. Nitrobacter acidophilus' ACIDNB-1 (WP_319798838.1)
Nitrobacter sp. Nb-311A (EAQ34794.1)
Nitrobacter sp. HKST-UBA79 (MCB1394038.1)
Nitrobacter sp. ECS31B4 (MCV0387649.1)
Chelativorans sp. AA-79 (WEX08601.1)
Hyphomicrobium denitrificans 1NES1 (AGK58303.1)
Methylocella tundrae T4 (VFU09812.1)
Pseudomonas luteola FDAARGOS_637 (QEU27571.1)
NarK2-like
NarK2-like
NarK1-like
NarK-c
Nitrococcus mobilis Nb-231 (EAR20979.1)
Escherichia coli str. K-12 substr. MG1655 (NP_415741.1)
Nitrospina watsonii (CAI2719027.1)
Nitrospina gracilis 3/211 (CCQ90797.1)
Nostoc punctiforme ATCC 29133 (B2IZT6)
Picosynechococcus sp. PCC 7002 (B1XLL7)
NrtP-like
Thermococcus litoralis DSM 5473 (NP_217330.1)
NrtP

**Supplementary Figure 8. Phylogenetic tree of nitrite/nitrate porter (NNP) family proteins.** A phylogenetic analysis of NNP family proteins, including NOB NarK proteins, reveals three distinct subgroups. NarK-n represents NarK proteins specific to *Nitrobacter*, NarK-c corresponds to *E. coli*-like NarK proteins (PDB-IDs: 4IU9 and 4U4V), and NrtP represents *Arabidopsis thaliana*–like high-affinity NNPs. NarK-n sequences that are associated with the *Nitrobacter* NXR gene cluster are highlighted in red. Branch support values of ≥ 95% are indicated by black circles. A complete list of NNP-family proteins from NOB included in the phylogenetic analysis is provided in **Supplementary Table 4**.

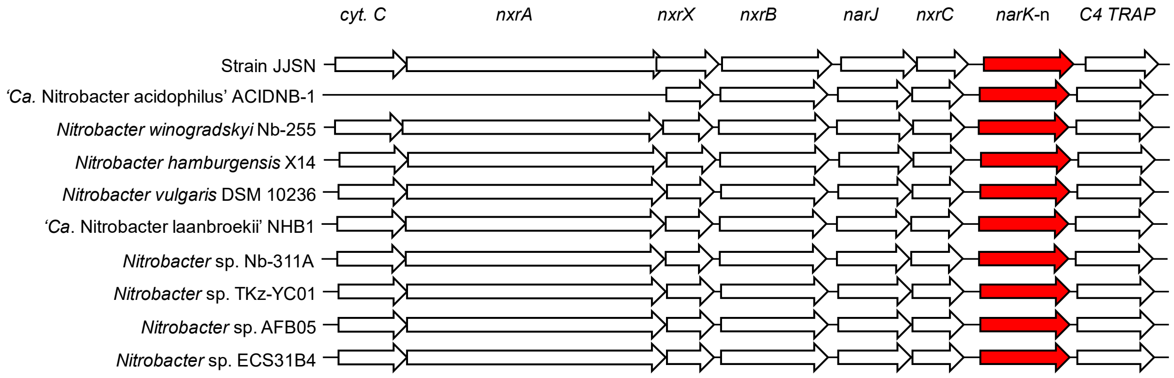


**Supplementary Figure 9. Conserved gene arrangement of the NXR cluster containing the *narK*-n gene in *Nitrobacter* strains.** Gene arrangements of *narK*-n (an NNP family nitrite/nitrate porter) and adjacent NXR-related genes are shown for selected Nitrobacter strains listed in **Supplementary Table 4**. All gene regions are oriented so that the NXR gene cluster is transcribed from left to right. *narK*-n is highlighted in red.

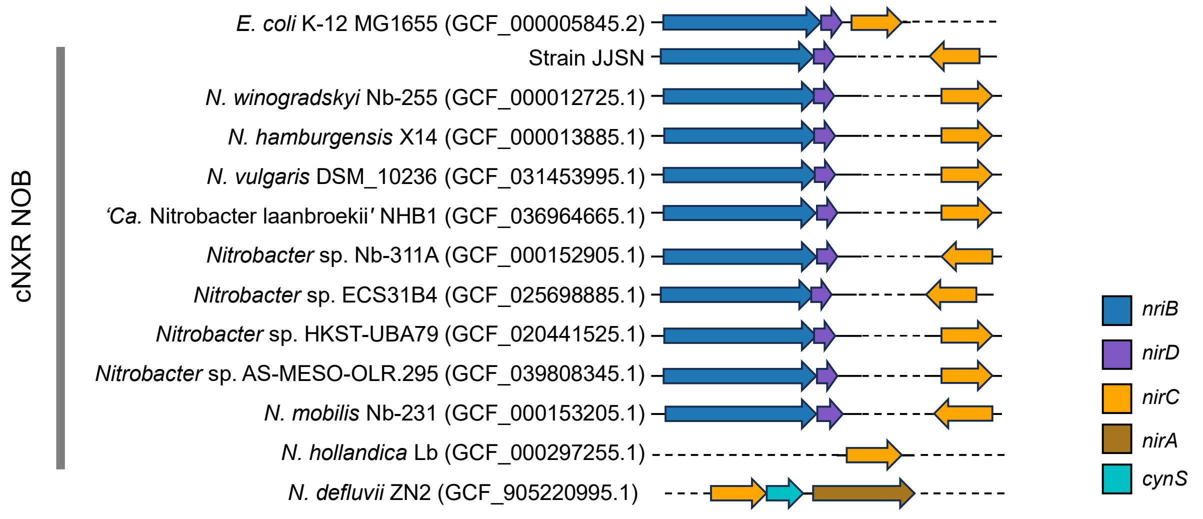


**Supplementary Figure 10. Genomic organization of the nitrite reductase (NIR)-associated gene clusters in NOB.** Genomic regions surrounding NIR-associated gene clusters are shown for *Escherichia coli* K-12 and selected NOB strains listed in **Supplementary Table 4**, including cNXR NOB (*Nitrobacter*, *Nitrococcus,* and *Nitrolancea*) and pNXR NOB (*Nitrospira defluvii* ZN2). Gene annotations are as follows: *nirB*, nitrite reductase (NADH) large subunit (blue); *nirD*, Nitrite reductase (NADH) small subunit (purple); *nirC*, formate/nitrite transporter family protein (orange); *nirA*, ferredoxin-nitrite reductase (brown); and *cynS*, cyanase (cyan). Genes depicted as arrows indicating the transcriptional direction. NCBI genome accession numbers are provided for each strain.

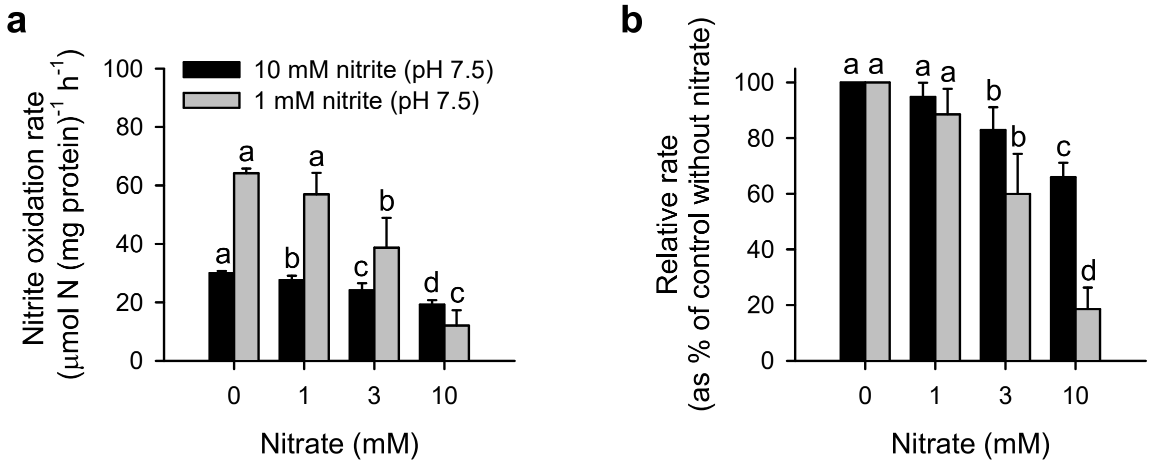


**Supplementary Figure 11. Effects of nitrate on the nitrite oxidation rate in *N. winogradskyi* Nb-255. a**, Absolute nitrite oxidation rates (μmol N mg protein$^{-1}$ h$^{-1}$) in cells grown at pH 7.5 with 10 mM nitrite (black) or 1 mM nitrite (gray), following exposure to varying nitrate concentrations as measured by microrespirometry. A substrate spike of 375 μM nitrite was used. **b**, Relative nitrite oxidation rates expressed as a percentage of the no-nitrate control (set to 100%). Bars represent the mean ± s.d. ($n$ = 3). Different letters represent significant differences ($p < 0.05$, one-way ANOVA followed by Tukey's test, performed separately for each nitrite concentration).

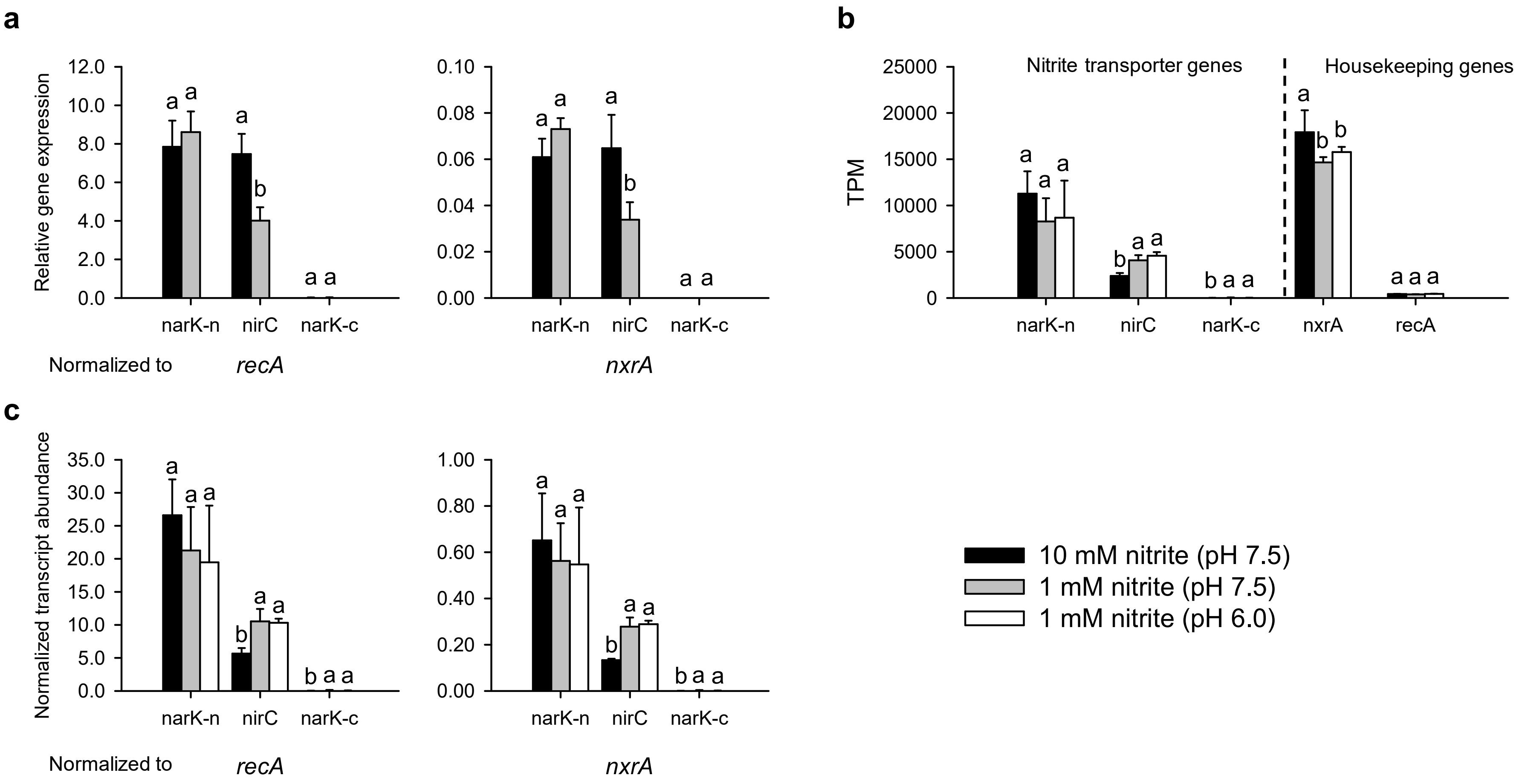
a
b
c
Relative gene expression
12.0
10.0
8.0
6.0
4.0
2.0
0.0
0.10
0.08
0.06
0.04
0.02
0.00
narK-n
nirC
narK-c
Normalized to
recA
nxrA
TPM
25000
20000
15000
10000
5000
0
Nitrite transporter genes
Housekeeping genes
nxrA
recA
Normalized transcript abundance
35.0
30.0
25.0
20.0
15.0
10.0
5.0
0.0
1.00
0.80
0.60
0.40
0.20
0.00
10 mM nitrite (pH 7.5)
1 mM nitrite (pH 7.5)
1 mM nitrite (pH 6.0)

**Supplementary Figure 12. Expression of nitrite transporter genes in *Nitrobacter winogradskyi* Nb-255 grown with 1 mM or 10 mM nitrite at pH 7.5 or 6.0. a,** Relative expression levels of *narK*-n, *nirC*, and *narK*-c were quantified by RT-qPCR and normalized to those of the housekeeping genes *recA* and *nxrA* (the latter being associated with the NXR gene cluster). Gene loci and primer information are provided in **Supplementary Table 6**. **b**, TPM values for nitrite transporter genes (*narK*-n, *narK*-c, *nirC*) and housekeeping genes (*recA* and *nxrA*) under three growth conditions are shown: 10 mM nitrite at pH 7.5 (black), 1 mM nitrite at pH 7.5 (gray), and 1 mM nitrite at pH 6.0 (white). **c**, Normalized transcript abundances of the same genes from transcriptomic data, also normalized to *recA* and *nxrA* (the latter associated with the NXR gene cluster). Bars represent mean ± s.d. ($n = 3$). Different letters denote statistically significant differences ($p < 0.05$, two-way ANOVA followed by Tukey's test). Raw transcriptome data are available in the **Supplementary Dataset 2.**

### Supplementary Dataset 1–3

**Supplementary Dataset 1: Genome annotations for strain JJSN.** This dataset provides the annotated genome of strain JJSN, including locus tags, gene names (if available), genomic coordinates, predicted protein products, and functional assignments based on multiple databases (e.g., COG, Pfam, TIGRfam, and KEGG Orthology)

**Supplementary Dataset 2: List of genes expressed in *N. winogradskyi* Nb-255 cells under three different growth conditions (*n* = 5):** 10 mM nitrite (pH 7.5); 1 mM nitrite (pH 7.5); 1 mM nitrite (pH 6.0). Expression categories indicate whether genes are upregulated (fold change > 1.5 and adjusted *p*-value < 0.05), downregulated (fold change < −1.5 and adjusted *p*-value < 0.05), or constitutively expressed (adjusted *p*-value > 0.05, or fold change between −1.5 and 1.5 regardless of significance).

**Supplementary Dataset 3: Expression data for nitrite transporter genes and housekeeping genes in *N. winogradskyi* Nb-255 (*n* = 15).** The data represent gene expression values under different nitrite concentrations and pH conditions from Supplementary Dataset 2. This dataset corresponds to Supplementary Figures 12b, c.
